# CRK2 and C-terminal phosphorylation of NADPH oxidase RBOHD regulate ROS production in Arabidopsis

**DOI:** 10.1101/618819

**Authors:** Sachie Kimura, Kerri Hunter, Lauri Vaahtera, Huy Cuong Tran, Matteo Citterico, Aleksia Vaattovaara, Anne Rokka, Sara Christina Stolze, Anne Harzen, Lena Meißner, Maya Melina Tabea Wilkens, Thorsten Hamann, Masatsugu Toyota, Hirofumi Nakagami, Michael Wrzaczek

## Abstract

Reactive oxygen species (ROS) are important messengers in eukaryotic organisms and their production is tightly controlled. Active extracellular ROS production by NADPH oxidases in plants is triggered by receptor-like protein kinase (RLK)-dependent signaling networks. Here we show that the cysteine-rich RLK CRK2 kinase activity is required for plant growth and CRK2 exists in a preformed complex with the NADPH oxidase RBOHD in Arabidopsis. Functional CRK2 is required for the full elicitor-induced ROS burst and consequently the *crk2* mutant is impaired in defense against the bacterial pathogen *Pseudomonas syringae pv*. tomato DC3000. Our work demonstrates that CRK2 regulates plant innate immunity. We identified *in vitro* CRK2-dependent phosphorylation sites in the C-terminal region of RBOHD. Phosphorylation of S703 RBOHD is enhanced upon flg22 treatment and substitution of S703 with alanine reduced ROS production in Arabidopsis. Phylogenetic analysis suggests that phospho-sites in C-terminal region of RBOHD are conserved throughout the plant lineage and between animals and plants. We propose that regulation of NADPH oxidase activity by phosphorylation of the C-terminal region might be an ancient mechanism and that CRK2 is an important element in regulating MAMP-triggered ROS production.

**One-sentence summary:** CRK2 associates with and activates RBOHD to trigger MAMP-induced ROS production and reveals a novel regulatory mechanism for plant NADPH oxidases through phosphorylation of the C-terminus.

## Introduction

Plants are continuously confronted with stimuli from the surrounding environment, including abiotic cues and invading pathogens. Plant cells also perceive a plethora of signals from neighboring cells and distant tissues. Numerous plasma membrane proteins are involved in the meticulous monitoring and transduction of signals for inter- and intracellular communication. A common early feature of many cellular responses to various environmental changes involves the production of reactive oxygen species (ROS) (Kimura et al., 2017; Waszczak et al., 2018). While ROS are an inevitable by-product of aerobic metabolism and their unrestricted accumulation can have deleterious consequences (Waszczak et al., 2018), ROS are also ubiquitous signaling molecules in plants and animals (Suzuki et al., 2011; Waszczak et al., 2018). Eukaryotic cells produce ROS in several subcellular compartments as well as the extracellular space, in plants referred to as apoplast (Kimura et al., 2017; Waszczak et al., 2018). A major component in the production of extracellular ROS is the evolutionarily conserved NADPH oxidase (NOX) family (Kimura et al., 2017; Meitzler et al., 2014). NOX-dependent ROS production is involved in regulation of immune functions, cell growth and apoptosis in animals and plants (Jiménez-Quesada et al., 2016; Waszczak et al., 2018).

Plant NOXs, referred to as respiratory burst oxidase homologs (RBOHs), have been identified as homologs of phagocyte gp91^*phox*^/NOX2, which contains six transmembrane helices and a C-terminal NADPH- and FAD-binding cytoplasmic region (Torres et al., 2002). Unlike gp91^*phox*^/NOX2, RBOHs contain an additional N-terminal region with Ca^2+^-binding EF-hands, similar to non-phagocytic NOXs, such as NOX5 (Suzuki et al., 2011). RBOH activity is strictly controlled to avoid damaging consequences of unrestricted ROS production (Suzuki et al., 2011). *Arabidopsis thaliana* (Arabidopsis) RBOHD is the best-characterized RBOH and is involved in biotic and abiotic stress responses (Lee et al., 2013; Lee et al., 2018; Torres et al., 2002). The N-terminal region of RBOHD is phosphorylated by a variety of protein kinases, including receptor-like cytoplasmic kinases (RLCKs) (Dubiella et al., 2013; Han et al., 2019; Kadota et al., 2014; Kaya et al., 2019; Kimura et al., 2012; Li et al., 2014; Ogasawara et al., 2008; Zhang et al., 2018), for example BOTRYTIS-INDUCED KINASE 1 (BIK1) (Kadota et al., 2014; Li et al., 2014). While previous research has suggested a predominant role of phosphorylation of the N-terminal region for regulation of RBOH, phosphorylation of the C-terminal region is important for the regulation of human gp91^*phox*^/NOX2 and NOX5 (Jagnandan et al., 2007; Raad et al., 2009). NADPH- and FAD-binding sites in C-terminus are highly conserved in NOXs and RBOHs, but it is unclear whether the C-terminus of plant RBOHs could also be a target for regulation of the ROS producing activity.

Apoplastic RBOH-dependent ROS production is a common response to the activation of receptor-like protein kinase (RLK; Shiu and Bleecker, 2001) signaling, in particular following perception of microbe-associated molecular patterns (MAMPs) or damage-associated molecular patterns (DAMPs; Couto and Zipfel, 2016; Kimura et al., 2017). However, the roles of the so-called ROS burst and its integration into RLK-triggered signaling networks are as yet unclear (Kimura et al., 2017). A large group of RLKs in plants is formed by the cysteine-rich RLKs (CRKs) (Vaattovaara et al., 2019). The extracellular region of CRKs harbors two copies of the domain of unknown function 26 (DUF26) but the molecular function of the CRK ectodomain remains unknown (Vaattovaara et al., 2019). CRKs have been linked to ROS signaling (Bourdais et al., 2015; Idänheimo et al., 2014; Yadeta et al., 2017; Yeh et al., 2015) and cell death (Bourdais et al., 2015; Yadeta et al., 2017). CRKs are important signaling elements in plant development, biotic and abiotic stress responses (Acharya et al., 2007; Bourdais et al., 2015; Chen et al., 2004; Chern et al., 2016; Hunter et al., 2019; Idänheimo et al., 2014; Tanaka et al., 2012; Wrzaczek et al., 2010; Yadeta et al., 2017; Yeh et al., 2015).

Here we characterize the role of CRK2 in immune signaling in response to MAMP-perception. CRK2 exists in a pre-formed complex with RBOHD. CRK2 controls the activity of RBOHD and functional CRK2 is required for full MAMP-induced ROS production. Importantly, we show that CRK2 phosphorylates the C-terminal region of RBOHD and modulates the ROS-production activity of RBOHD *in vivo.* Our results lead us to propose a novel mechanism for the regulation of RBOHD activity through phosphorylation of the C-terminal region and highlight a critical role for CRK2 in the precise control of the ROS burst in response to biotic stress.

## Results

### CRK2 kinase activity is important for plant development

CRK2 has been previously implicated in stress responses and development in Arabidopsis (Bourdais et al., 2015). CRK2 is a typical CRK with N-terminal signal peptide, extracellular region containing two DUF26 domains, transmembrane region and intracellular protein kinase domain (Figure 1A). The *crk2* mutant was smaller than wild type (Col-0) plants (Figure 1B) (Bourdais et al., 2015), and displayed significantly reduced fresh (Figure 1C) and dry weight (Figure S1A). Over-accumulation of the plant hormone salicylic acid (SA) often leads to a reduction of plant size, but SA levels were not significantly different between *crk2* and wild type plants (Figure S1B). Expression of YFP-tagged CRK2 under the control of the CRK2 promoter *(CRK2pro::CRK2-YFP)* in *crk2* background restored plant growth (Figures. 1B, 1C and S1A). Substitution of the ATP-binding lysine (K) at position 353 with glutamic acid (E; CRK2^K353E^) or the aspartic acid (D) at position 450 in the catalytic domain VIb (Stone and Walker, 1995) with asparagine (N; CRK2^D450N^) abated the kinase activity of CRK2 *in vitro* (Hunter et al., 2019). Kinase dead CRK2^K353E^-YFP or CRK2^D450N^-YFP under control of the *CRK2* promoter expressed in leaves and roots, and displayed the same subcellular localization as wild type CRK2-YFP (Figures S1C and S1D) but failed to restore the growth defect of *crk2* (Figures 1B, 1C and S1A). The amino acid substitutions did not reduce the abundance of kinase dead CRK2-YFP (Figure S1E). In summary, our results show that CRK2 is important for proper plant growth and its kinase activity is crucial for this function.

**Figure 1.**
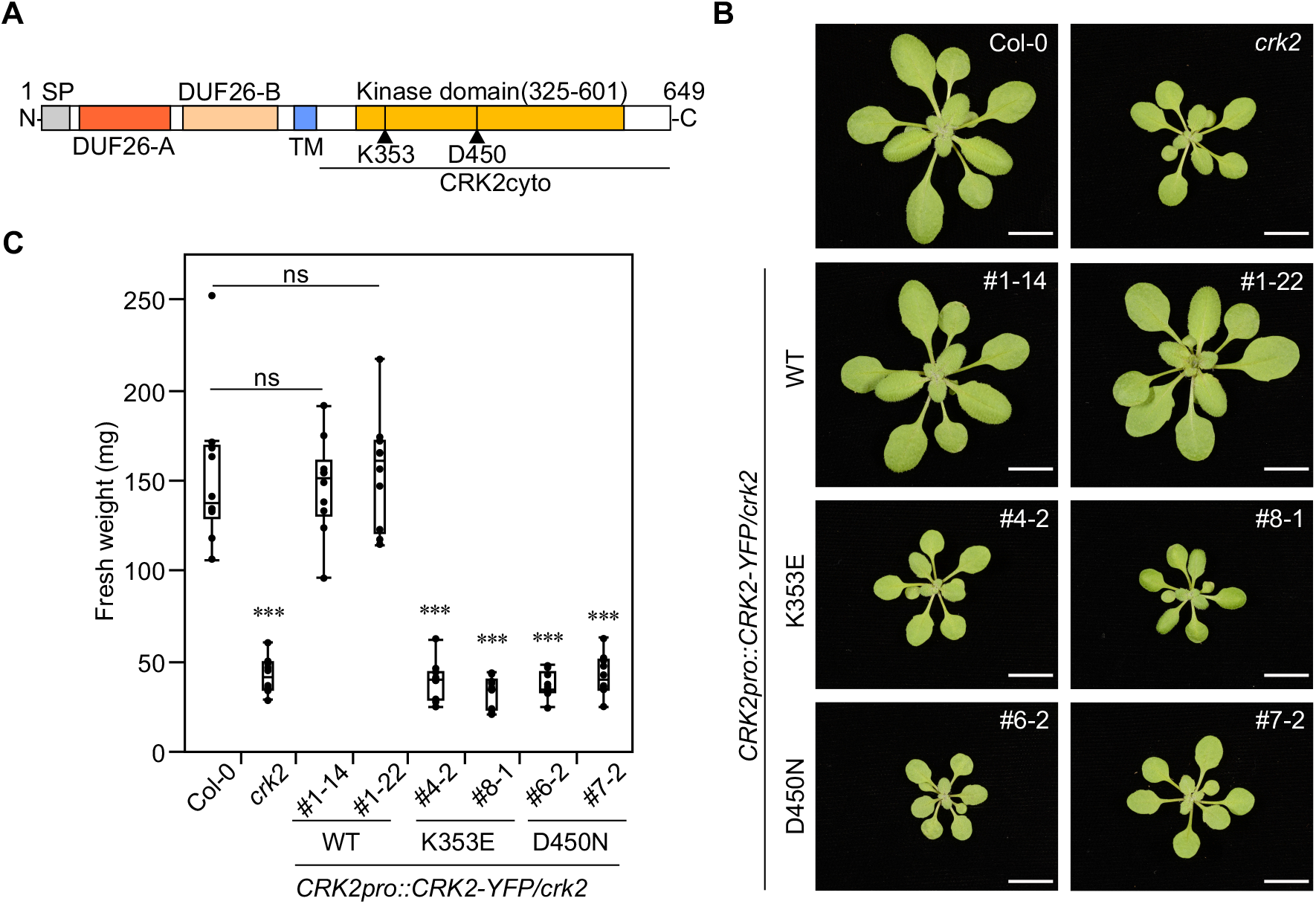
CRK2 kinase activity is required for plant growth. **(A)** Schematic representation of CRK2 structure. SP: signal peptide (AAs 1-29), DUF26-A (AAs 39-132), DUF26-B (AAs 146-243), TM: transmembrane domain (AAs 261-283), and kinase domain (AAs 325-601). **(B)** Representative pictures of 21-day-old plants of Col-0, *crk2, CRK2pro::CRK2-YFP/crk2, CRK2pro::CRK2^K353E^-YFP/crk2* and *CRK2pro::CRK2D^450N^-YFP/crk2* plants. Bar = 1 cm. **(C)** Box plot shows the fresh weight of 21-day-old plants (n = 10). Differences between Col-0 and transgenic lines were evaluated with One-way Anova with Tukey-Kramer HSD, *** p<0.001, ns, not statistically significant. The experiment was repeated three times with similar results.

### CRK2 is required for MAMP-triggered responses and resistance to *Pseudomonas syringae* pv. tomato DC3000

Previous results suggested that ROS production triggered by flg22, a MAMP derived from bacterial flagella, is reduced in *crk2* (Bourdais et al., 2015). Therefore, we tested the role of CRK2 in MAMP-induced ROS production in detail. ROS production triggered by flg22 was reduced in *crk2* and reintroduction of CRK2-YFP into the mutant background restored ROS production to the same levels as in Col-0 (Figure 2A). The flg22-induced ROS production in plants expressing CRK2^D450N^-YFP was comparable to *crk2* (Figure 2B). Transcriptional upregulation of flg22 responsive genes *(FRK1* and *NHL10*) showed that MAMP-perception was not impaired in *crk2* (Figures S2A and S2B). To test whether the reduced response of *crk2* to flg22 was accompanied by altered pathogen susceptibility, we measured growth of the hemibiotrophic bacterial pathogen *Pseudomonas syringae* DC3000 pv. tomato (*P*to DC3000). The *crk2* mutant was significantly more susceptible to the virulent pathogen compared to Col-0 (Figure 2C). CRK2-YFP but not the kinase-dead CRK2^D450N^-YFP rescued the pathogen susceptibility of *crk2* (Figure 2C). ROS production induced by chitin (Figure S2C) or pep1 (Figure S2D) was also reduced in *crk2* compared to Col-0 suggesting that the reduced MAMP- or DAMP-triggered ROS production in *crk2* is a general response and not specific to flg22. MAMP-triggered ROS production is also an important element in pathogen-triggered stomatal closure. Therefore, we tested whether the reduced flg22-triggered ROS production in *crk2* also coincided with altered stomatal closure. The *crk2* mutant displayed slightly reduced stomatal aperture compared to wild type plants under control conditions. However, treatment with flg22 triggered stomatal closure in wild type but not in *crk2* plants (Figure S2E).

**Figure 2.**
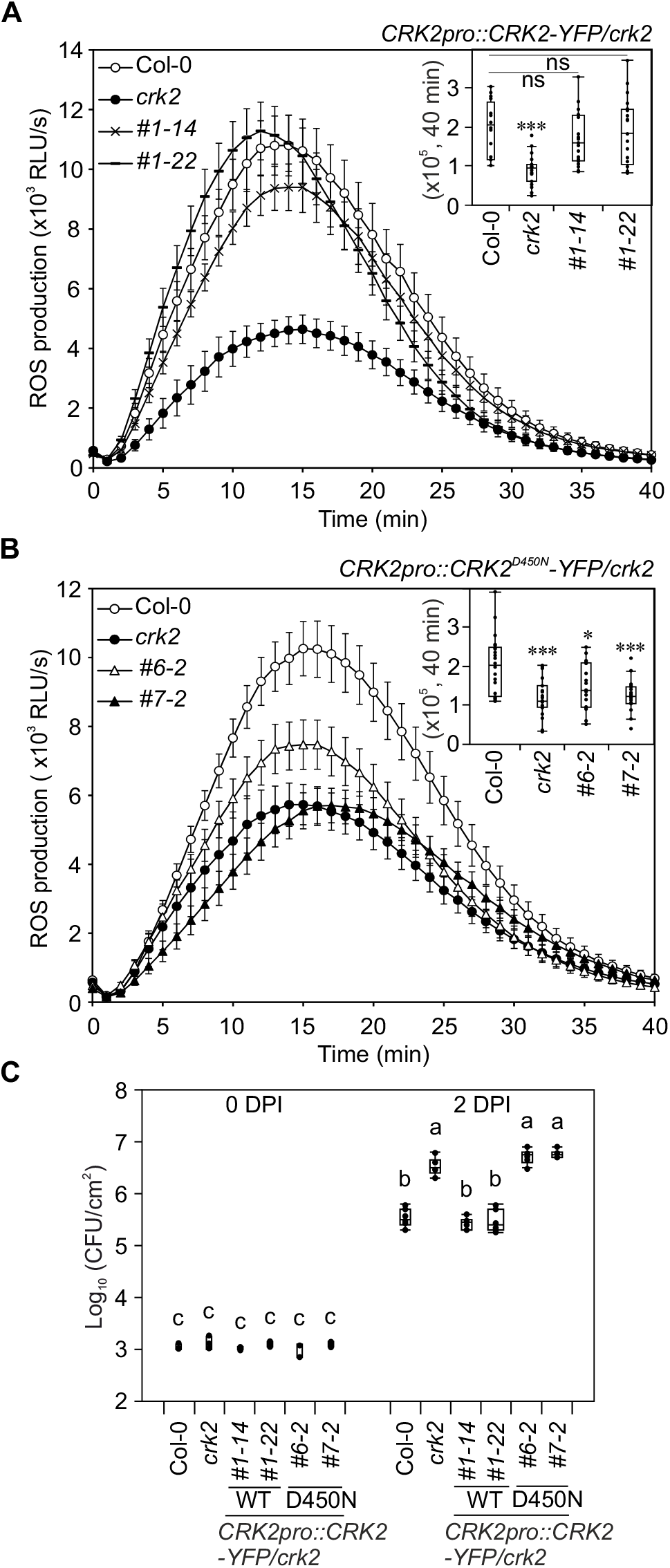
CRK2 regulates flg22-triggered ROS production and resistance to a virulent bacterial pathogen. **(A)** and **(B)** flg22-induced ROS production in Col-0, *crk2* and *CRK2pro::CRK2-YFP/crk2* or *CRK2pro::CRK2D^450N^-YFP/crk2.* Leaf discs from 28-day-old plants were treated with 200 nM flg22 and ROS production was measured. Box plot shows cumulative ROS production over 40 min (upper right). **(A)** Values represent mean ±SEM of n ≤ 16. Differences compared with Col-0 were evaluated with One-way Anova with Tukey-Kramer HSD, *** p < 0.001, ns, not statistically significant. **(B)** Values represent the mean ±SEM of n ≤ 19. Differences compared with Col-0 were evaluated with Oneway Anova with Tukey-Kramer HSD, *p < 0.05, *** p < 0.001. **(C)** Quantitative analysis of bacterial growth in Col-0, *crk2* and *CRK2pro::CRK2-YFP/crk2 or CRK2pro::CRK2D^450N^-YFP/crk2* following syringe infiltration with *Pto* DC3000 (1 x 10^5^ CFU/mL). Box plot shows the numbers of bacteria in leaf discs after infiltration [0 day post infiltration (0 DPI), n = 3] or 2 DPI (n = 6). Each data point contains 4 leaf discs. Letters indicate significant differences at p < 0.05 (One-way Anova with Tukey-Kramer HSD). **(A)** to **(C)** The experiment was repeated three times with similar results.

To further investigate the role of CRK2 in flg22-triggered responses, we assessed Ca^2+^ signaling, MAPK activation and callose deposition in *crk2.* Application of flg22 resulted in a rapid increase in cytosolic Ca^2+^ ([Ca^2+^]_cyto_) levels in wild type plants, which express the FRET-based Ca^2+^-sensor *YCNano-65* (Choi et al., 2014; Lenglet et al., 2017; Toyota et al., 2018). This response was reduced in *YCNano65/crk2* background, (Figures S2F and S2G). Interestingly, flg22-dependent MAPK activation (Figure S2H) was more pronounced in *crk2* compared to Col-0. Callose deposition was enhanced in *crk2* compared to Col-0 after 30 minutes of treatment with flg22 (Figures S2I and S2J). However, flg22-triggered callose deposition was similar in Col-0 and *crk2* after 12 hours (Figure S2I). Taken together, CRK2 is an essential component for mounting immune responses against *P* to DC3000 in Arabidopsis, modulating extracellular ROS production, stomatal closure, Ca^2+^ influx, MAPK activation and early callose deposition.

### CRK2 interacts with RBOHD and controls ROS production

RBOHD is the main source of MAMP/DAMP-induced extracellular ROS production (Couto and Zipfel, 2016; Kimura et al., 2017) and flg22-, pep1 and chitin-induced ROS production was significantly reduced in *crk2* (Figures 2 and S2). RBOH proteins, including RBOHD, are synergistically activated by protein phosphorylation and Ca^2+^-binding to EF-hand motifs in the N-terminal region (Kaya et al., 2019). Given the importance of CRK2 kinase activity in MAMP-induced ROS production we investigated whether CRK2 could activate RBOHD. To test this, we used human embryonic kidney 293T (HEK293T) cells, a human cell culture which produces minimal amounts of extracellular ROS due to a lack of expression of endogenous NADPH oxidases (Ogasawara et al., 2008). HEK293T cells were co-transfected with *3FLAG-RBOHD* and *CRK2-3Myc* or *3Myc-GFP* as control. Subsequently, RBOHD-mediated extracellular ROS production was measured by luminol-amplified chemiluminescence. Despite equal 3FLAG-RBOHD protein levels (Figure S3A) ROS production in cells co-transfected with *CRK2-3Myc* and *3FLAG-RBOHD* was strongly elevated compared to cells co-transfected with *3FLAG-RBOHD* and *3Myc-GFP* (Figure 3A). Co-transfection with the inactive variant *CRK2^D450N^-3Myc* did not enhance ROS production by *3FLAG-RBOHD* compared to cotransfection with *CRK2-3Myc* (Figure 3A). Transfection of *CRK2-3Myc* in the absence of *3FLAG-RBOHD* did not induce ROS production in HEK293T cells (Figure 3A). Since RBOHD can also be activated by Ca^2+^, HEK293T cells were treated with ionomycin, a Ca^2+^ ionophore that induces a rise in cytosolic Ca^2+^ levels. Ionomycin-induced transient ROS production (Δ_delta_ ROS: ROS_T=30_ to ROS_T=31_) in *CRK2-3Myc* and *3FLAG-RBOHD* co-transfected cells was not different from *3Myc-GFP* and *3FLAG-RBOHD* co-transfected cells (Figure 3A). Enhancement of RBOHD activity by CRK2 was not dependent on Ca^2+^ influx as the elevated basal ROS production activity of RBOHD co-transfected with *CRK2-3Myc* (ROS_T=0_ to ROS_T=30_ in Figure 3A) was also observed when using Ca^2+^-free assay buffer (Figures S3B and S3C). These results suggest that *CRK2-3Myc* enhanced the basal ROS-producing activity of 3FLAG-RBOHD in HEK293T cells uncoupling it from Ca^2+^ dependence.

**Figure 3.**
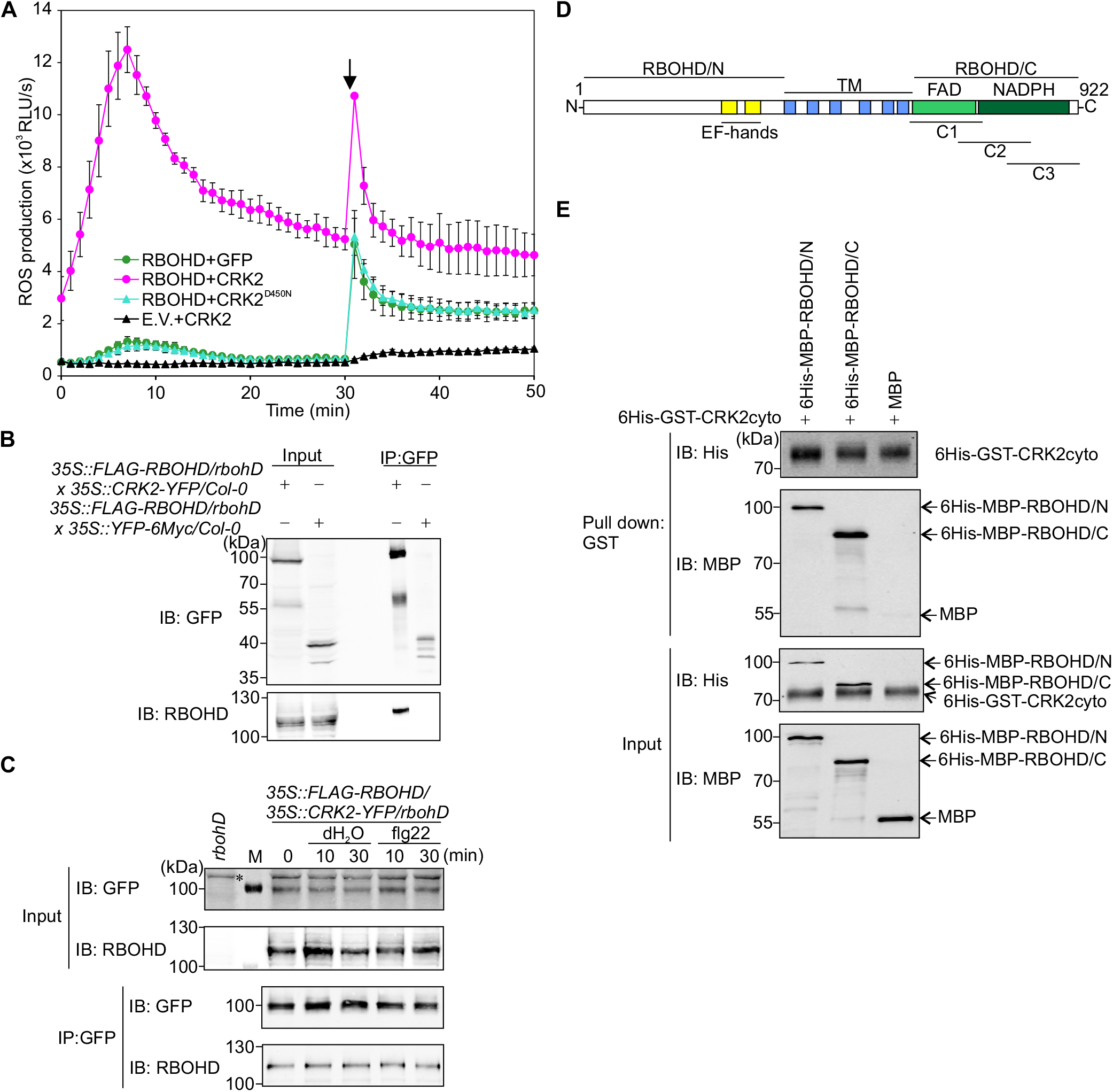
CRK2 interacts with RBOHD. **(A)** ROS production of RBOHD-expressing HEK293T cells. 3FLAG-RBOHD was transiently co-expressed with either 3Myc-GFP or CRK2 (WT or D450N)-3Myc. After 30 min 1 μM ionomycin was added to the medium (black arrow). Values represent mean ±SEM of n = 3. E.V. = empty vector. The experiment was repeated three times with similar results. **(B)** and **(C)** Co-IP analysis of interaction between RBOHD and CRK2. CRK2-YFP was immunoprecipitated using anti-GFP beads followed by immunoblotting with anti-RBOHD and anti-GFP antibodies. FLAG-RBOHD: 105 kDa, CRK2-YFP: 99.9 kDa and YFP-6Myc: 36.7 kDa. **(B)** *35S::FLAG-RBOHD/rbohD* × *35S::CRK2-YFP/Col-0* (F1) and *35S::FLAG-RBOHD/rbohD × 35S::YFP-6Myc/Col-0* (F1) plants. The experiment was repeated three times with similar results. **(C)** *35S::FLAG-RBOHD/35S::CRK2-YFP/rbohD* plants with 1 μM flg22 treatment. M: Protein molecular marker, *: unspecific signal. Total protein from *rbohD* was used for immunoblot of input as a negative control. **(D)** Schematic representation of RBOHD structure. EF-hands (AAs 257-329), TM: transmembrane domains (AAs 374 - 605), FAD: FAD-binding domain (AAs 613-730), NADPH: NADPH-binding domain (AAs 736-904), RBOHD/N: RBOHD N-terminal region (AAs 1-376), RBOHD/C: RBOHD C-terminal region (AAs 606-922); C1: RBOHD/C1 (AAs 606-741), C2: RBOHD/C2 (AAs 696-831), C3: RBOHD/C3 (AAs 787-922). **(E)** *In vitro* pull-down analysis of direct interaction between RBOHD and CRK2. MBP, 6His-MBP-RBOHD/N and 6His-MBP-RBOHD/C were incubated with 6His-GST-CRK2cyto and pull down with GST followed by immunoblotting with anti-6His and anti-MBP antibodies. 6His-GST-CRK2cyto: 68.5 kDa, 6His-MBP-RBOHD/N: 84.7 kDa, 6His-MBP-RBOHD/C: 78.4 kDa, MBP: 50.8 kDa. The experiment was repeated two times with similar results.

The Arabidopsis genome encodes 10 *RBOHs* (Kaya et al., 2019). To test whether CRK2 specifically activates RBOHD, *CRK2-3Myc* was co-transfected with *3FLAG-RBOHF* and *3FLAG-RBOHC* into HEK293T cells. CRK2-3Myc enhanced basal ROS-producing activity of RBOHC and RBOHF in HEK293T cells similarly to RBOHD (Figures S3D-S3G). However, while the basal ROS production activity (ROS_T=5_) of RBOHD and RBOHF was elevated approximately 10-fold, the basal activity of RBOHC was only elevated 3-fold.

To investigate whether CRK2 and RBOHD interact *in planta,* we performed co-immunoprecipitation (Co-IP) assays using *rbohD* plants expressing *35S::CRK2-YFP* and *35S::FLAG-RBOHD*. CRK2-YFP was immunoprecipitated using an anti-GFP antibody coupled to magnetic beads and co-purified RBOHD was detected using a RBOHD-specific antibody. RBOHD co-purified with CRK2 (Figure 3B) and treatment of plants with flg22 did not alter the interaction of CRK2 with RBOHD (Figure 3C). To analyze this interaction in more detail, we carried out *in vitro* interaction assays between the cytosolic region of CRK2 (Figure 1A) and the cytosolic N-terminal and C-terminal regions of RBOHD (Figure 3D). Recombinant RBOHD/N or RBOHD/C tagged with 6His and maltose-binding protein (MBP; 6His-MBP-RBOHD/N, 6His-MBP-RBOHD/C) or MBP were incubated with the cytosolic region of CRK2, which contains the kinase domain (CRK2cyto), tagged with 6His and glutathione S-transferase (GST; 6His-GST-CRK2cyto) and glutathione sepharose beads. GST pull-down assay showed that 6His-GST-CRK2cyto interacted *in vitro* with 6His-MBP-RBOHD/N but intriguingly also with 6His-MBP-RBOHD/C (Figure 3E). In summary, our results suggest that CRK2 is capable of direct interaction with RBOHD. CRK2 and RBOHD form a complex, which exists independent of flg22 perception *in planta,* in contrast to many other RLK-containing complexes which are formed in response to ligand-binding.

### CRK2 phosphorylates RBOHD *in vitro*

The kinase activity of CRK2 was essential for the full flg22-triggered ROS burst *in planta* as well as for enhancing ROS production by RBOHD in HEK293T cells. Therefore, we tested whether CRK2 could phosphorylate RBOHD *in vitro.* Recombinant 6His-GST-CRK2cyto and 6His-MBP tagged RBOHD cytosolic regions (Figures 1A and 3D) were produced in *E. coli* and affinity purified. The 6His-GST-CRK2cyto phosphorylated 6His-MBP-RBOHD/N but not MBP (Figure 4A). Because of the similar molecular weight of 6His-GST-CRK2cyto (68.5 kDa) and 6His-MBP-RBOHD/C (78.4 kDa), RBOHD/C was divided into three overlapping fragments (C1, C2, and C3; Figure 3D). The results showed that the C1 and C3 fragments of 6His-MBP-RBOHD were preferentially phosphorylated by 6His-GST-CRK2cyto while C2 displayed considerably weaker phosphorylation (Figure 4B). Mass spectrometric analysis of in-gel trypsin- or Lys-C-digested peptides identified *in vitro* RBOHD phosphorylation sites targeted by CRK2cyto (Table S1). In the N-terminal region of RBOHD, two sites targeted by CRK2 were identified (S8 and S39), while three sites (S611, S703, S862) were identified in the C-terminal region. Taken together our results show that the N- and C-terminal regions of RBOHD are phosphorylated by CRK2 *in vitro.*

**Figure 4.**
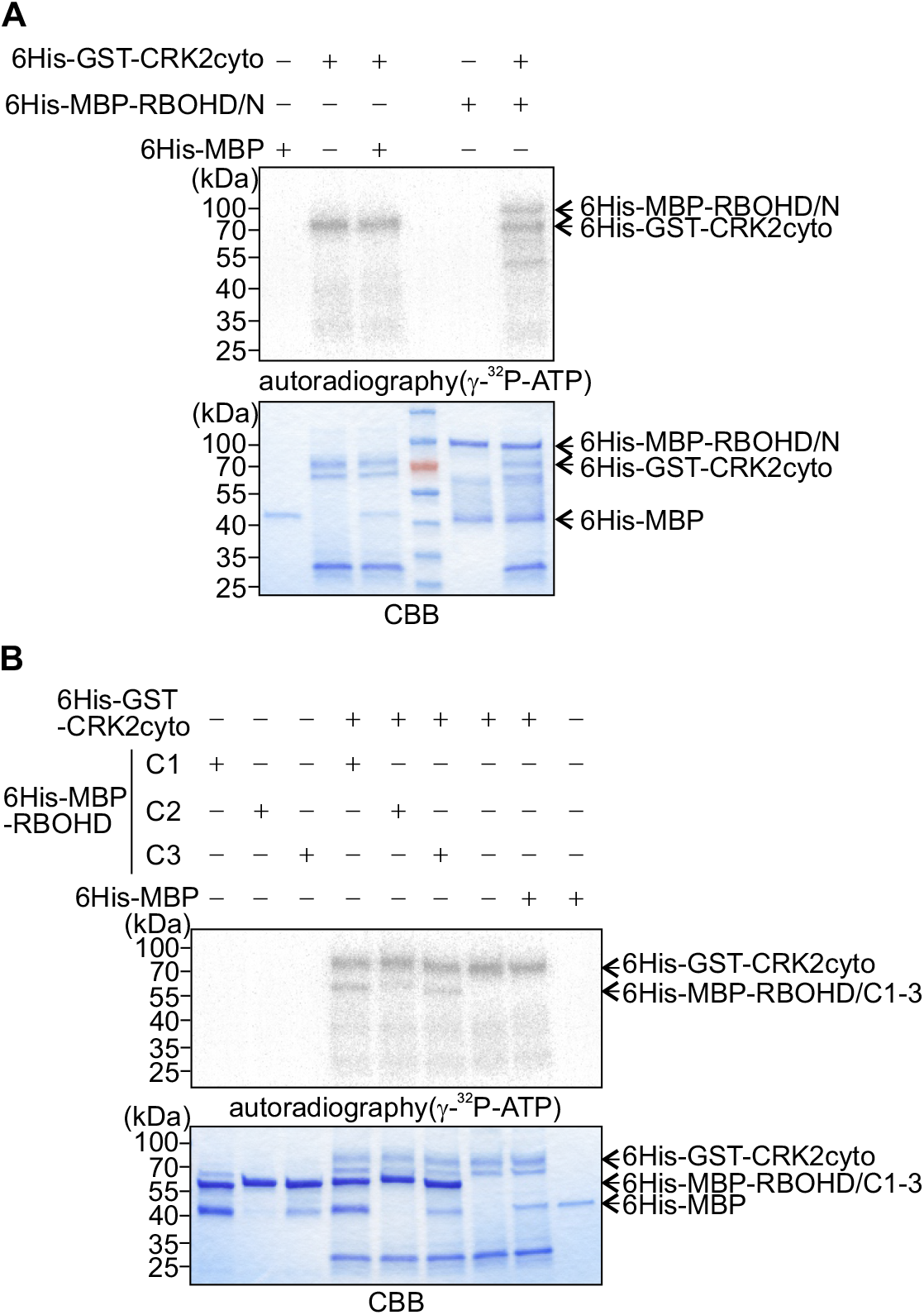
CRK2 phosphorylates the cytosolic regions of RBOHD *in vitro.* **(A)** and **(B)** Autophosphorylation and transphosphorylation were visualized with [γ-^32^P] ATP and autoradiography (upper panel). Input proteins were stained with coomassie brilliant blue (CBB) (lower panel). Experiments were repeated three times with similar results. 6His-GST-CRK2cyto: 68.5 kDa, 6His-MBP-RBOHD/N: 84.7 kDa, 6His-MBP-RBOHD/C1:57.9 kDa, /C2:57.8 kDa, /C3:58.4 kDa, 6His-MBP: 44.3 kDa. **(A)** *In vitro* transphosphorylation of 6His-MBP-RBOHD N-terminus by 6His-GST-CRK2cyto. 6His-MBP-RBOHD/N or 6His-MBP was incubated with 6His-GST-CRK2cyto in kinase buffer. **(B)** *In vitro* transphosphorylation of 6His-MBP-RBOHD C-terminus by 6His-GST-CRK2cyto. 6His-MBP-RBOHD/C1, /C2, /C3 or 6His-MBP was incubated with 6His-GST-CRK2cyto in kinase buffer.

### CRK2 regulates RBOHD *via* phosphorylation of S703 and S862

Among the RBOHD phospho-sites targeted by CRK2 *in vitro,* phosphorylation of S8 and S39 has been previously described to be phosphorylated by SIK1 (Zhang et al., 2018) and BIK1 (Kadota et al., 2014; Li et al., 2014). S703 has been reported to be phosphorylated upon xylanase treatment but no responsible kinase was identified (Benschop et al., 2007) while phosphorylation of S611 and S862 has not been described so far. In order to test whether the identified phospho-sites in RBOHD were important for the regulation of RBOHD activity, the residues S8, S39, S611, S703, and S862 were substituted with alanine to make them non-phosphorylatable. Wild type RBOHD and phospho-site mutant constructs were transfected into HEK293T cells together with CRK2. Amino acid substitutions did not affect RBOHD protein levels (Figures S4A and S4B). Substitution of S8 or S39 in the N-terminal cytoplasmic region of RBOHD did not impact ROS-producing activity compared to the wild type protein when co-transfected with CRK2 (Figure 5A). The 3FLAG-RBOHD^S703A^ and CRK2-3Myc co-transfected cells showed reduced basal ROS production as compared to 3FLAG-RBOHD and CRK2-3Myc, (Figure 5B), suggesting that S703 could be a positive regulatory site for RBOHD activity. In contrast to 3FLAG-RBOHD^S703A^, HEK293T cells expressing 3FLAG-RBOHD^S862A^ and CRK2-3Myc exhibited higher basal ROS production compared to 3FLAG-RBOHD and CRK2-3Myc (Figure 5B), suggesting that S862 could act as a negative regulatory site. ROS production of 3FLAG-RBOHD^S611A^ co-transfected with CRK2-3Myc was similar to 3FLAG-RBOHD suggesting no regulatory role of this single site. Mutation of S703 or S862 of RBOHD did not impair Camdependent activation of ROS production of RBOHD itself without CRK2 co-transfection (Figures S4D and S4D). Taken together, our results suggest that the phospho-sites in the C-terminal cytoplasmic region of RBOHD could be crucial for fine-tuning ROS production activity in HEK293T cells.

**Figure 5.**
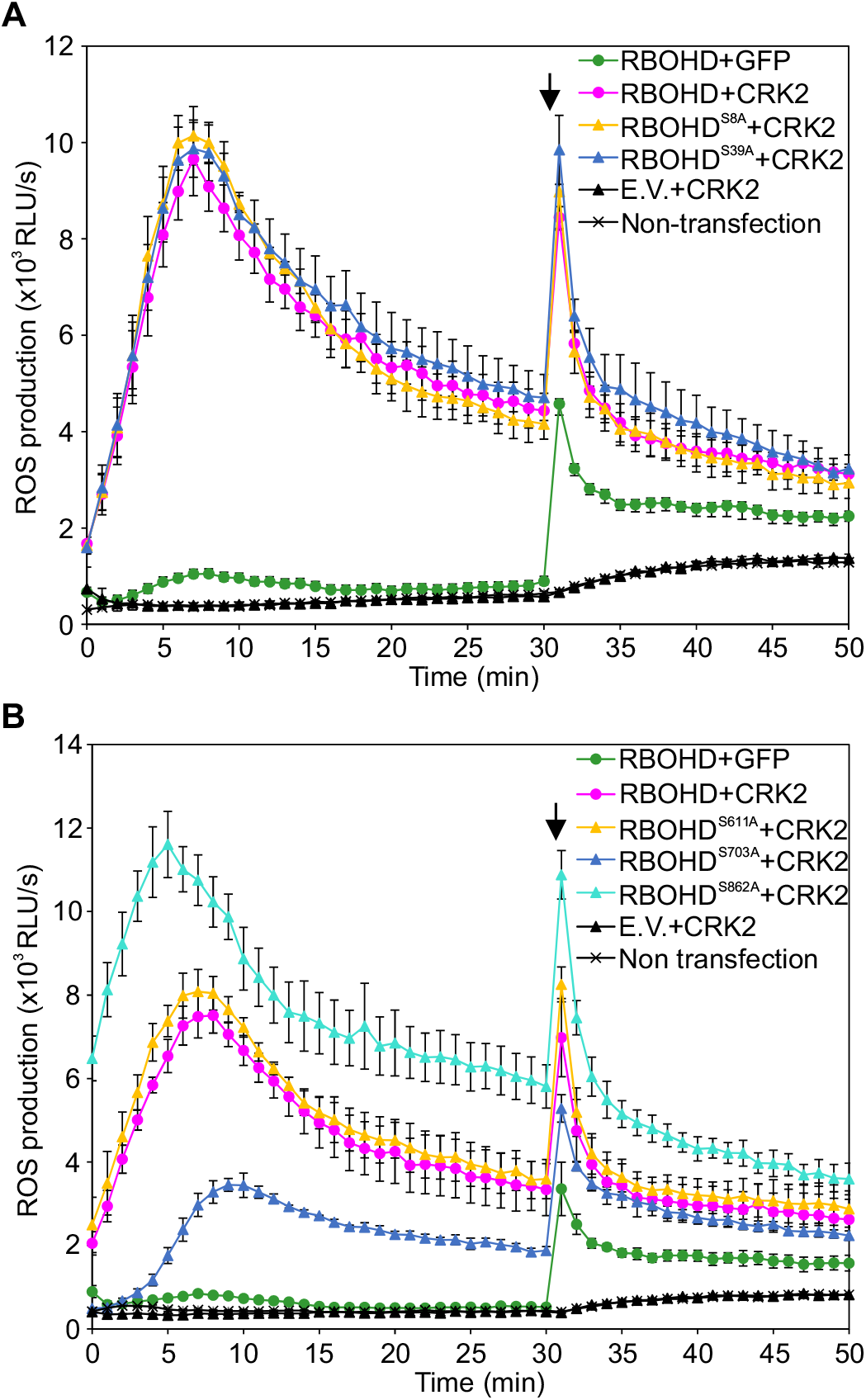
CRK2 modulates the ROS-production activity of RBOHD *via* phosphorylation of the C-terminus in HEK293T cells. **(A)** and **(B)** Effect of mutations of CRK2-dependent *in vitro* phosphorylation sites in the N-terminal **(A)** or C-terminal **(B)** cytosolic region of RBOHD. After 30 min 1 μM ionomycin was added to the medium to promote Ca^2+^ influx (black arrow). Values represent mean ±SEM of n = 3. E.V. = empty vector. The experiment was repeated three times with similar results. **(A)** 3FLAG-RBOHD (WT, S8A or S39A) was transiently co-expressed with either 3Myc-GFP or CRK2-3Myc in HEK293T cells. **(B)** 3FLAG-RBOHD (WT, S611A, S703A or S862A) were transiently co-expressed with either 3Myc-GFP or CRK2-3Myc in HEK293T cells.

### Phosphorylation of S703 of RBOHD modulates flg22-induced ROS production *in planta*

To investigate whether RBOHD phosphorylation sites targeted by CRK2 *in vitro* were also phosphorylated upon flg22-treatment *in planta*, we carried out targeted phosphoproteomic analyses of Col-0 seedlings treated with flg22 for 5 min. Phosphorylation of S8 was not significantly induced by flg22-treatment (Figures 6A, S5A and S5B) while S39 phosphorylation was strongly enhanced (Figures 6B and S5C). Phosphorylation of S611 and S862 could not be evaluated as trypsin or Lys-C digestion resulted in phosphosite-containing peptides of inappropriate length for LC-MS-based targeted analyses. However, phosphorylation of S703, which was targeted by CRK2 *in vitro,* and mutation of which to alanine reduced ROS production in HEK293T cells, was significantly enhanced upon flg22 treatment (Figures 6C and S5D). In agreement with previous studies, phosphorylation of S163, S343, and S347 in RBOHD (Figures S5E-S5G), as well as dual phosphorylation of the TEY-motif in the MAPKs MPK3, MPK6 and MPK11 (Figures S5H-S5J), were enhanced by flg22-treatment (Kadota et al., 2015). To test whether phosphorylation of S703 in the C-terminal region of RBOHD was mediated by CRK2 in *planta,* we used leaf discs of *CRK2pro::CRK2 (WT or D450N)-YFP/crk2.* The basal and flg22-induced phosphorylation of S8 and S39 were not significantly altered in *CRK2pro::CRK2-YFP/crk2* or the kinase dead variant (Figures S5K-S5M), suggesting that CRK2 might not be the main kinase phosphorylating RBOHD S8 and S39 *in planta.* However flg22-induced phosphorylation of S703 in RBOHD was significantly reduced in kinase dead *CRK2pro::CRK2(D450N)-YFP/crk2* plants compared *CRK2pro::CRK2-YFP/crk2* (Figure 6D). These results suggest that CRK2 is responsible, at least partially, for flg22-induced phosphorylation of S703 in *planta.*

**Figure 6.**
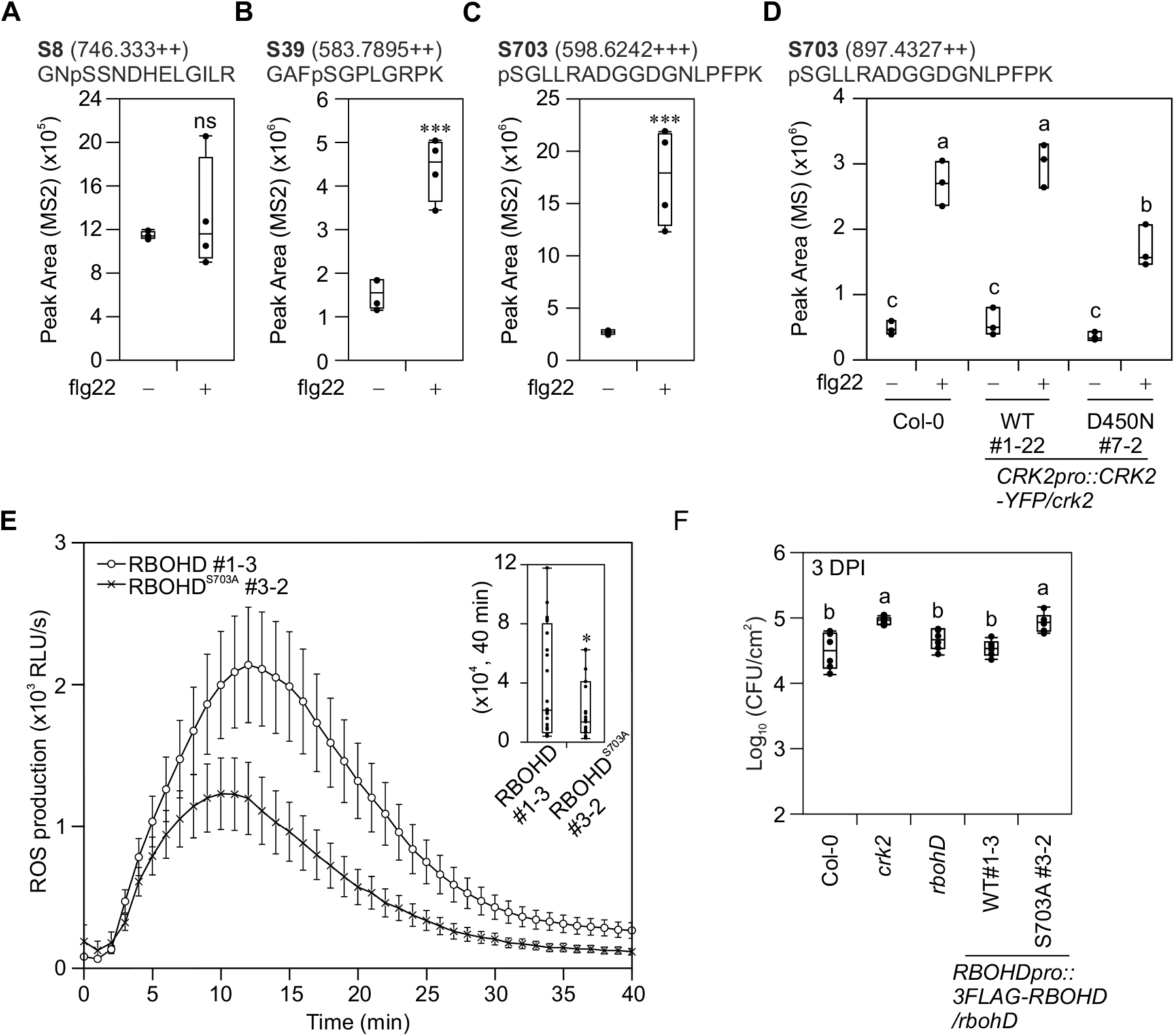
RBOHD S703 is involved in regulation of flg22-induced ROS production. **(A)** to **(C)** Quantification of RBOHD phosphorylation in Col-0 upon flg22 treatment. 12-day-old seedlings were treated with water (-) or 1 μM flg22 (+) for 5 min. Total proteins were digested with trypsin (S8 and S39) or Lys-C (S703) and phosphopeptides were enriched, and then selected phosphopeptides were quantified by LC-MS/MS. Box plots show MS2 fragment peak ion areas of indicated phosphopeptides (n = 4). Differences between water- or flg22-treated samples were evaluated with One-way Anova with Tukey-Kramer HSD, *** p<0.001, ns, not statistically significant. **(A)** RBOHD S8 residue. **(B)** RBOHD S39 residue. **(C)** RBOHD S703 residue. **(D)** Quantification of RBOHD S703 phosphorylation in Col-0, *CRK2pro::CRK2-YFP/crk2* #1-22 and *CRK2pro::CRK2D^450N^-YFP/crk2* #7-2 upon flg22 treatment. Leaf discs from 28-day-old plants were treated with water (-) or 200 nM flg22 (+) for 5 min. Total proteins were digested with Lys-C and phosphopeptides were enriched, and then selected phosphopeptides were quantified by LC-MS/MS. Box plots show MS2 fragment peak ion areas of indicated phosphopeptides (n = 3). Each data point contains 14 leaf discs. Letters indicate significant differences at p < 0.05 (One-way Anova with Tukey-Kramer HSD). **(E)** flg22-induced ROS production in *RBOHDpro::3FLAG-RBOHD/rbohD* #1-3 and *RBOHDpro::3FLAG-RBOHD^S703A^/rbohD* #3-2. Leaf discs from 28-day-old plants were treated with 200 nM flg22. Box plot shows cumulative ROS production over 40 min (upper right). Values represent mean ±SEM of n ≥ 23. Difference between lines was evaluated with One-way Anova with Tukey-Kramer HSD, * p < 0.05. The experiment was repeated three times with similar results. **(F)** Quantitative analysis of bacterial growth in Col-0, *crk2, rbohD, RBOHDpro::3FLAG-RBOHD/rbohD* #1-3 and *RBOHDpro::3FLAG-RBOHD^S703A^/rbohD* #3-2 following spray inoculation with *Pto* DC3000 (1 x 10^5^ CFU/mL). Box plot shows the numbers of bacteria in leaf discs 3 days post inoculation (3 DPI) (n = 6). Each data point contains 4 leaf discs. Letters indicate significant differences at p < 0.05 (One-way Anova with Tukey-Kramer HSD).

To investigate whether phosphorylation of S703 in the C-terminal region of RBOHD also impacts RBOHD-dependent ROS production *in planta,* we generated transgenic plants expressing RBOHD or RBOHD^S703A^ under the control of the *RBOHD promoter (RBOHDpro::3FLAG-RBOHD)* in *rbohD* background. The phospho-site mutations did not alter growth or development compared to the 3FLAG-RBOHD expressing plants (Figure S6A). Protein amounts of 3FLAG-RBOHD or 3FLAG-RBOHD^S703A^ were comparable in the transgenic plant lines (Figures S6B and S6C). ROS production triggered by flg22 was significantly reduced in 3FLAG-RBOHD^S703A^ plants compared to 3FLAG-RBOHD plants (Figures 6E and S6D). The results suggest that phosphorylation of S703 in the C-terminus of RBOHD is important for full flg22-triggered ROS production also *in planta.* Notably, expression of 3FLAG-RBOHD^S703A^ in *rbohD* mutant background impaired flg22-triggered stomatal closure (Figure S6E) and consequently enhanced susceptibility to *P*toDC3000 after spray infection (Figure 6F) and. In summary, our results suggest phosphorylation of RBOHD S703 is an important regulatory element of plant immune responses.

### C-terminal phosphorylation sites are conserved in plant and animal NADPH oxidases

Since little is known about control of RBOH activity *via* its C-terminus we investigated whether regulation through S703 was unique to RBOHD or conserved also in other RBOHs. We constructed a phylogenetic tree of RBOHs from plant genomes representing major branches of the plant lineages (Figure 7). Plant RBOHs form a monophyletic group, which is separated from human NOX2 and NOX5. The C-terminal region of plant NADPH oxidases contains a high number of serines and threonines (Table S2). Furthermore, conservation of the amino acid sequence surrounding a phosphorylation site may be required for recognition of the site by the kinase (Miller and Turk, 2016; Trost et al., 2016). Accordingly, the sites corresponding to RBOHD S611, S703 and S862 exhibited conservation of amino acid motifs surrounding the phosphorylation site.

**Figure 7.**
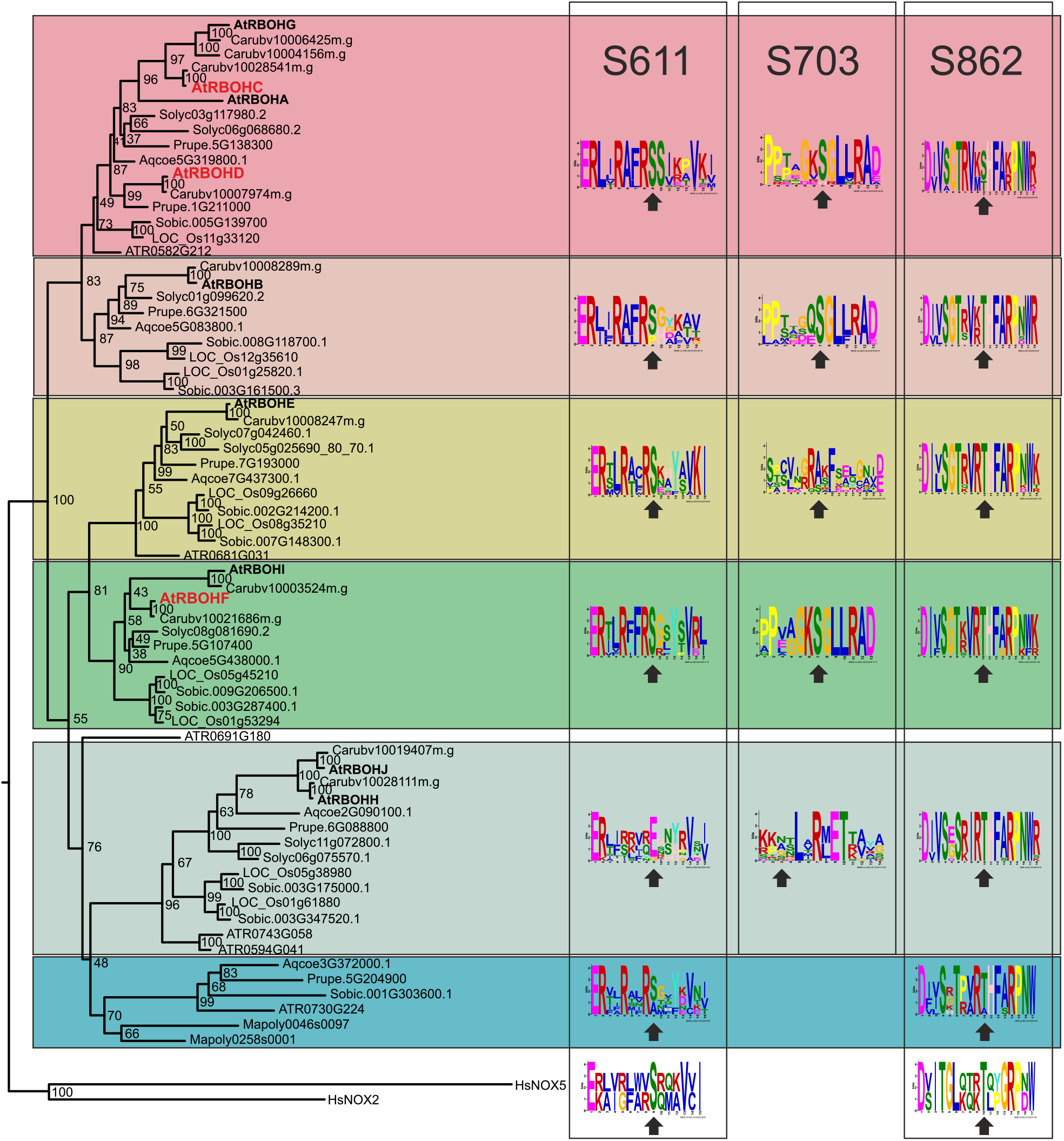
Phosphorylation sites in the C-terminal region are conserved in plants and animals. Phylogenetic tree showing that plant RBOHs form a single clade which is parallel to the NADPH oxidases NOX2 and NOX5ß from *Homo sapiens.* The tree was constructed using RAxML from a MAFFT alignment, 1000 bootstraps were calculated with RAxML. The full sequence alignment can be found in Wasabi at http://was.bi?id=p_nwi1. Plant species included were: *Arabidopsis thaliana* (At), *Capsella rubella* (Carub), *Prunus persica* (Prupe), *Solanum lycopersicum* (Solyc), *Aquilegia coerulea* (Aqcoe), *Oryza sativa* (LOC_Os), *Sorghum bicolor* (Sobic), *Amborella trichopoda* (Atr), and *Marchantia polymorpha* (Mapoly). Numbers of phospho-sites in the meme figures represent the position of the amino acid in RBOHD from *Arabidopsis thaliana.* Arrows indicate the position of the phospho-site (S or T) or corresponding amino acid.

The phospho-sites in the C-terminal region displayed strong conservation throughout the plant RBOH clade. S703 was conserved in eight of the ten RBOHs from Arabidopsis but not in RBOHH and RBOHJ (Figure 7). In the clade containing AtRBOHE the sequence motif harboring S703 was far less conserved compared to other clades. Intriguingly, AtRBOHE itself as well as its homolog from the *Brassicaceae Capsella rubella* contains a serine corresponding to S703 of RBOHD while the other members of the clade do not possess a serine or threonine at this position. The phospho-sites S862, which may be involved in negative regulation based on experiments in HEK293T cells (Figure 5B), as well as S611 were strongly conserved in all plant RBOHs. Intriguingly, the sequence motifs harboring S611 and S862 are conserved even in human NOX2 and NOX5 (Figure 7). The C-terminal region binds FAD and NADPH. Therefore, it may underlie strong evolutionary constraints to conserve these binding properties. This may be also reflected in the strong conservation of C-terminal phospho-sites and their sequence context in plant and animal NADPH oxidases alike.

## Discussion

CRKs are a large group of RLKs involved in biotic and abiotic stress signaling in Arabidopsis (Bourdais et al., 2015). We have previously shown that flg22-triggered extracellular ROS production is altered in several *crk* mutants (Bourdais et al., 2015). In particular CRK2, a member of the basal clade of CRKs (Vaattovaara et al., 2019), has been highlighted since *crk2* displays striking phenotypes (Bourdais et al., 2015) including reduced rosette size and diminished flg22-induced ROS production. Functional CRK2 restored the reduced rosette size (Figure 1) as well as the MAMP-triggered ROS burst (Figure 2). In addition to their roles in stress responses, extracellular ROS have also been implicated in leaf cell expansion (Schmidt et al., 2016). For example, the *rbohD rbohF* double mutant displays reduced rosette size (Torres et al., 2002). Overexpression of CRKs has been associated with increased SA accumulation (Acharya et al., 2007; Chen et al., 2004), which could potentially explain the growth phenotype of *crk2.* However, since SA levels were unaltered in the loss-of-function mutant *crk2* (Figure S1B), its smaller size may rather be a consequence of impaired ROS production (Figure 2). This is supported by the observation that CRK2 enhanced the activity of RBOHD and RBOHF in HEK293T cells (Figures 3A and S3F). Alternatively, other proteins interacting with or substrates of CRK2 might be involved in the regulation of plant growth.

In line with reduced MAMP- and DAMP-induced ROS production (Figures 2A, 2B, S2C and S2D), *crk2* was more susceptible to the virulent bacterial pathogen *P*to DC3000 (Figure 2C) suggesting that CRK2-mediated ROS production was essential to effectively counteract pathogen infection. Also other flg22-induced defense responses were altered in *crk2* including reduced changes in cytosolic Ca^2+^ and reduced stomatal closure and MAPK activation (Figures S2E-S2H). Ca^2+^ is an important element in the activation of RBOH. Moreover, ROS also triggers Ca^2+^ fluxes in plants and contributes to stomatal closure (Kimura et al., 2017). Thus, the diminished increase of cytosolic Ca^2+^ and stomatal closure in *crk2* may be a consequence of the impaired flg22-induced ROS production. Also, callose deposition (Caillaud et al., 2014; Ellinger et al., 2014) has been previously linked to ROS production (Couto and Zipfel, 2016) and CRK2 interacts with callose synthases, phosphorylates CALLOSE SYNTHASE 1 (CALS1) *in vitro* and salt-induced callose deposition is reduced in *crk2* (Hunter et al., 2019). While flg22-triggered callose deposition in *crk2* was indisthinguable from Col-0 after 12 hours of treatment (Figure S2I and J), *crk2* exhibited more callose deposition after 30 minute treatment with flg22 compared to Col-0. Interestingly, CRK2 forms clusters at the plasma membrane in response to flg22-treatment and ROS is required for this process (Hunter et al., 2019). It is not clear how these clusters are integrated with the regulation of RBOHD activity but it might serve to connect RBOHD-dependent ROS production with callose deposition. Another important element in response to biotic and abiotic cues is the activation of MAPK cascades (Bigeard et al., 2015; Boudsocq et al., 2015) and earlier reports suggest a bifurcation to ROS burst and MAPK activation in defense signaling following MAMP-perception (Yeh et al., 2016; Zhang et al., 2007). CRK2 could be involved in balancing MAMP-induced ROS signaling pathways and MAPK signaling but the mechanisms are still unclear.

Our results suggest that CRK2 participates in the control of ROS production *via* interaction with RBOHD. Intriguingly, CRK2 existed in a pre-formed complex with RBOHD *in planta* independent of MAMP-treatment (Figure 3C) while many other RLK protein complexes are formed upon signal perception. The observation that, similar to flg22-triggered ROS production, also chitin-induced ROS production was reduced in *crk2* suggests that CRK2 might not act on MAMP receptor complexes but rather on the regulation of ROS production itself. Phosphorylation of the C-terminus is critical for the regulation of human NADPH oxidases. Phosphorylation of the NOX2 C-terminus by protein kinase C (PKC) enhances assembly of the multimeric NOX2 complex and its activity, whereas phosphorylation by *ataxia telangiectasia-mutated* (ATM) kinase inhibits NOX2 activity (Beaumel et al., 2017; Raad et al., 2009). NOX5 activity is regulated by Ca^2+^-binding to EF-hands in the N-terminus (Banfi et al., 2004) but NOX5 is also activated by phosphorylation of the C-terminus by PKCα or calcium/calmodulin-dependent kinase II (CAMKII; Chen et al., 2014; Pandey et al., 2011). Although the C-terminal catalytic domain of RBOHs is highly conserved in plants and animals, the N-terminus has been considered as important for activation of the ROS-production activity. Accordingly, multiple phospho-sites (S8, S39, S133, S148, S163, S339, S343 and S347) in the N-terminal region have been reported. Intriguingly, CRK2cyto interacted with and phosphorylated the RBOHD C-terminal region at S611, S703 and S862 (Figures 3E, 4B and Table S1). Phosphorylation of S703 upon xylanase treatment has been reported (Benschop et al., 2007) but not linked with other MAMPs or modulation of ROS production. Treatment with flg22 enhanced phosphorylation of S703 in Arabidopsis (Figures 6C, 6D and S5D). Mutation S703A in RBOHD led to reduced CRK2-dependent RBOHD activity in HEK293T cells (Figure 5B) and decreased flg22-induced ROS production in Arabidopsis (Figures 6E and S6D). The flg22-induced phosphorylation of S703 was diminished in *CRK2^D450N^-YFP/crk2* plants compared to wild type (Figure 6D). These results suggest that CRK2-mediated phosphorylation of the RBOHD C-terminus at S703 contributes to the regulation of MAMP-induced ROS production. Phosphorylation sites in the C-terminus are highly conserved among RBOHs (Figure 7) suggesting that phosphorylation of the C-terminal region could be a general feature of plant NADPH oxidases. Remarkably, two putative RBOHD phospho-sites, S611 and S862, were identified even in the human NADPH oxidases NOX2 and NOX5 (Figure 7).

Substitution of RBOHD S862 to alanine resulted in enhanced ROS-producing activity in HEK293T cells. However, substitution of RBOHD S611 to alanine, similarly to RBOHD S39A, did not alter ROS production in HEK293T cells (Figure 5). RBOHD S39A also did not affect flg22-induced ROS production but phosphomimic S39D enhanced the ROS production and phosphorylation of S39 is enhanced by MAMP treatment *in planta* (Kadota et al., 2014) (Figures 6B and S5C). These results suggest the importance to determine the phosphorylation status of S611 and S862 *in planta* in the future. RBOHD can also be regulated by cysteine S-nitrosylation in the C-terminus (Yun et al., 2011) but it is unclear how this modification is integrated with other regulatory mechanisms. Taken together, our results suggest that phosphorylation of the C-terminal region of plant NADPH oxidases is strongly conserved and important for controlling ROS production.

Several protein kinases phosphorylate RBOHD N-terminus and regulate the activity including RLCKs (Kadota et al., 2014; Li et al., 2014; Lin et al., 2015), MAP4Ks (Zhang et al., 2018), CPKs (Dubiella et al., 2013) and RLKs (Chen et al., 2017) but how is regulation by phosphorylation of the N- and C-terminal regions coordinated? BIK1 activates RBOHD by phosphorylation (Kadota et al., 2014; Li et al., 2014) and ROS production in *bik1* is reduced to a similar extent as in *crk2.* However, decreased flg22-induced ROS production in *crk2* was not due to lower *BIK1* transcript abundance (Figure S7). BIK1 homologs, AvrPphB SUSCEPTIBLE1 (PBS1) and AvrPphB SUSCEPTIBLE1-LIKE (PBL) kinases, contribute to the regulation of RBOH activity and ROS production is progressively reduced in double mutants of *bik1* with its homologs (Lin et al., 2015; Zhang et al., 2018). CRK2 and BIK1 might synergistically regulate ROS production but we were unable to obtain a double mutant between *bik1* and *crk2* (Table S3). Therefore, we propose that at least one of these components is essentially required. BIK1 has previously been shown to interact with other kinases including CRKs (Lee et al., 2017) but interaction with CRK2 has not been investigated. BIK1 and CRK2 are likely highly coordinated in order to precisely control ROS production in response to environmental stimuli (Figure 8). Like CRKs, RBOHs are involved in diverse processes in stress responses and also plant development and it is conceivable that different CRKs regulate the diverse set of RBOH proteins in various cellular contexts potentially *via* phosphorylation of the C-terminal region.

**Figure 8.**
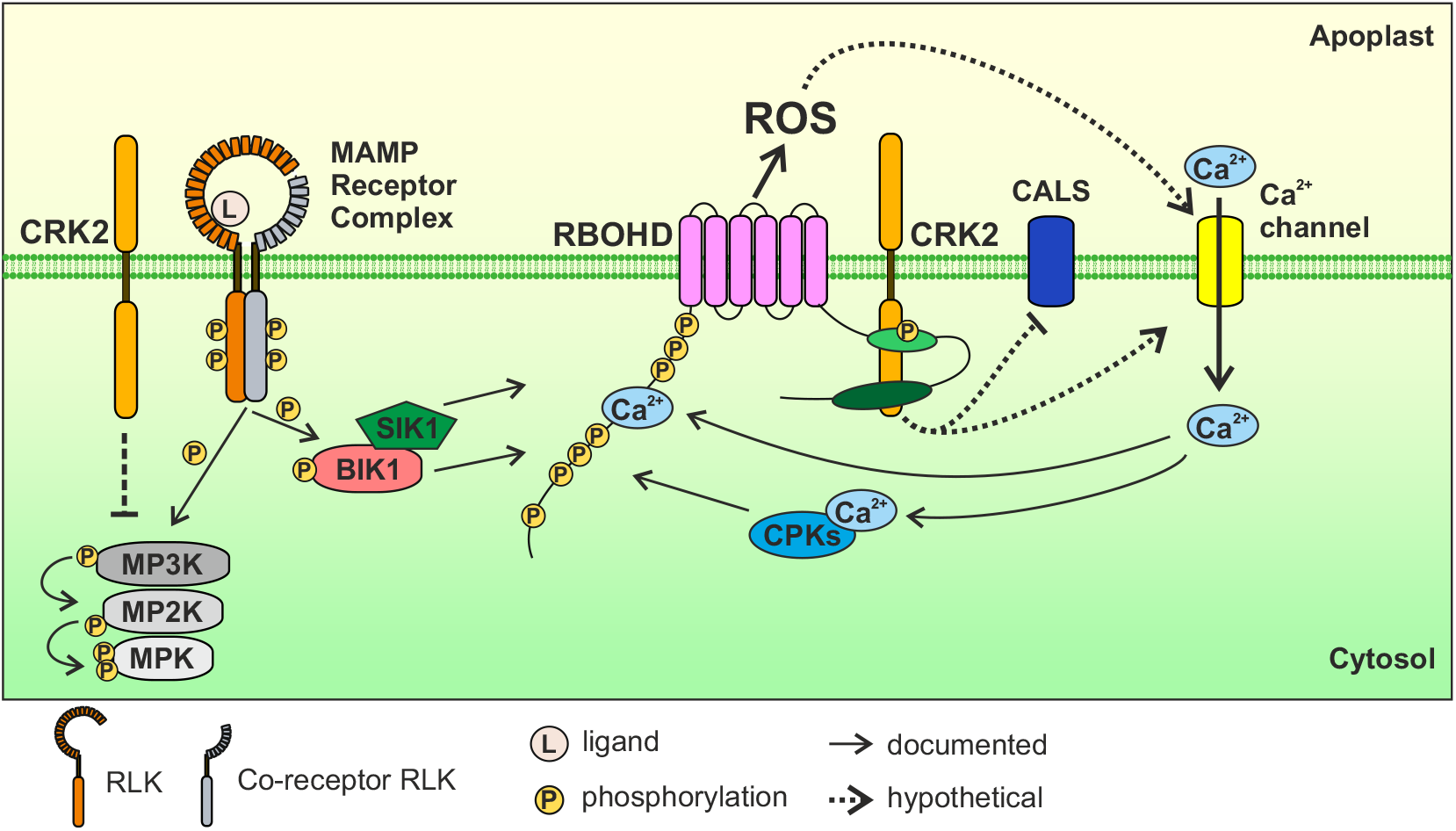
Schematic model for MAMP-triggered RBOHD activation. MAMPs are recognized by MAMP receptor complexes. RBOHD N-terminus is phosphorylated by BIK1 and SIK1 and apoplastic ROS production is induced. Apoplastic ROS production by RBOHD leads to Ca^2+^ influx into the cytosol. Ca^2+^-binding to RBOHD N-terminus and to CPKs leads to Ca^2+^-dependent activation of RBOHD. We found that CRK2 also contributes to the activation of RBOHD *via* phosphorylation of its C-terminus at S703. CRK2 can also mediates inhibition of MAPK activation and callose deposition *via* CALS after MAMP perception. MPK, mitogen-activated protein kinase; MP2K, MPKK; MP3K, MPKKK.

In summary, we propose that CRK2 is a central element in orchestrating the extracellular ROS burst and in mediating the balance between different defense responses. The full complexity and integration of the regulatory components controlling RBOH activity is still a topic of much speculation (Kimura et al., 2017). The diversity of regulators converging at RBOHs reflects the prominent role of apoplastic ROS in signal transduction while strict control of their activity is required to circumnavigate oxidative damage. We suggest that RBOHD is regulated by phosphorylation of the C-terminal region to complement regulatory mechanisms targeting the N-terminus (Figure 8). Based on the conservation of amino acid motifs in the C-terminus of NADPH oxidases harboring serine and threonine residues (Figure 7) we propose that this mode of regulation could be evolutionarily conserved in plants and animals. In the future it will be interesting to investigate how CRK-mediated phosphorylation of the RBOH C-terminus is integrated in the diverse processes which incorporate extracellular ROS.

## Materials and methods

### Plant Material and growth condition

*Arabidopsis thaliana* plants used in this study include Col-0, *crk2* (Bourdais et al., 2015), *rbohD* (Torres et al., 2002), *fls2* (Zipfel et al., 2004), *bik1* (Veronese et al., 2006), *35S::FLAG-RBOHD/rbohD* (Kadota et al., 2014), *35S::CRK2-YFP/Col-0* and *35S::YFP-6Myc/Col-0* (Hunter et al., 2019). To generate *crk2/bik1* double mutant, *crk2* and *bik1* single mutant plants were crossed. F1, F2 and F3 progenies were analyzed by PCR. F2 and F3 seeds were obtained by self-pollination. Primers are listed in Table S4.

Seeds were sterilized by 70 % ethanol 2 % Triton X-100 for 5 min and washed 3 times with 99 % ethanol. Surface sterilized seeds were sown on 1× or ½ strength Murashige and Skoog (MS) medium containing 1 % sucrose and subsequently stratified for 2-4 days in the dark at 4 °C. Plants were grown in growth chambers (Panasonic, #MLR-352-PE) under 12 h light/12 h dark (22 °C /18 °C) at 145 μmol m^-2^ s^-1^ (Panasonic FL40SS ENW/37). After 10 days, seedlings were transferred to soil and grown in growth rooms under the following conditions: 12 h light/12 h dark (23 °C /19 °C) at 220-280 μmol m^-2^ s^-1^ (Osram L 36W/865 Lumilux Cool Daylight or Osram L 58W/865 Lumilux Cool Daylight), relative humidity 50-60 %, unless otherwise stated.

For SA quantification, seeds were sown in commercial soil with perlite (Hasselfors Garden Yrkeskvalitet S-jord). Seeds were stratified for 3 days at 4 °C, then grown for a week in long-day conditions (16 h light, 22 °C 40 % humidity / 8 h dark, 18 °C, 60 % humidity) in 150 μmol m^-2^ s^-1^ photon flux density with a light source Philips Master TL5 HE 21W/830. Seedlings were then transplanted into fresh soil, and grown for 18 days more before sampling.

### Cell culture and Transfection

HEK293T cells (ATCC, #CRL-3216) were maintained at 37 °C in 5 % CO2 in Dulbecco’s Modified Eagle’s Medium nutrient mixture Ham’s F-12 (SIGMA, #D8062) supplemented with 10 % fetal bovine serum (Gibco, #26140-079). Cells were transfected with *pcDNA3.1* and *pEF1* vectors using GeneJuice transfection regent (Merck Millipore, #70967-3) according to the manufacturer’s instructions.

### Plasmid construction

*CRK2* and *RBOHD* constructs for Arabidopsis were generated through MultiSite Gateway technology (Invitrogen). To generate pBm43GW-CRK2pro::CRK2-Venus (YFP)-3AT for *crk2* complementation lines, the coding region of *CRK2* or kinase-dead mutants (K353E or D450N) were recombined into pENTR/D-TOPO vector (Invitrogen). pDONRP4P1R/zeo-CRK2pro, pDONR/zeo-CRK2 (or pENTR/D-TOPO-CRK2 kinase-dead mutant) and p2R3a-VenusYFP-3AT were recombined with pBm43GW. To generate pHm43GW-pRBOHD::3FLAG-RBOHD-nosT, the coding region of 3FLAG-RBOHD was amplified by PCR from pcDNA3.1-3FLAG-RBOHD and cloned into pDONR/zeo vector (Invitrogen). The promoter region of *RBOHD* was amplified by PCR from pBin19g-pRBOHD::3FLAG-RBOHD and cloned into pDONRP4P1R/zeo vector (Invitrogen). pDONRP4P1R/zeo-RBOHDpro, pDONR/zeo-3FLAG-RBOHD and p2R3a-nosT were recombined with pHm43GW. Single amino acid substitution mutants of *CRK2* and *RBOHD* were generated by point-mutant primers and the mega-primer PCR method. pBm43GW-CRK2pro::CRK2-YFP-3AT and pHm43GW-pRBOHD::3FLAG-RBOHD-nosT constructs were transformed into *crk2* and *rbohD* plants, respectively, by *Agrobacterium tumefaciens* strain GV3101 (pSoup)-mediated floral dipping (Clough and Bent, 1998). p2R3a-Venus(YFP)-3AT, p2R3a-nosT, pBm43GW and pHm43GW (Siligato et al., 2016), pBin19g-pRBOHD::3FLAG-RBOHD (Kadota et al., 2014), pcDNA3.1-3FLAG-RBOHD (Kaya et al., 2019), pDONR/zeo-CRK2 and pDONRP4P1R/zeo-CRK2pro (Hunter et al., 2019) have been described previously.

6His-GST-CRK2cyto and 6His-MBP-RBOHD/C constructs for recombinant proteins were generated by using In-Fusion technology (Clontech). The coding regions of CRK2cyto (WT, K353E, and D450N), RBOHD/C (full-length, C1, C2, and C3) were amplified by PCR and cloned into pOPINK (Addgene, #41143) or pOPINM (Addgene, #26044) vectors. pOPINM-RBOHD/N was described previously (Kadota et al., 2014).

For HEK293T cell experiments, pEF1-MCS-3Myc [*Bam*HI-*Not*I-3Myc-stop fragment was inserted between *KpnI* and *XbaI* sites of pEF1/myc-His B vector (Invitrogen)] was generated. To generate pEF1-CRK2 (WT or D450N)-3Myc, the codon optimized coding sequence of Kozak-CRK2 (WT or D450N) was cloned between *BamHI* and *NotI* sites of pEF1-MCS-3Myc. To generate pcDNA3.1-3FLAG-RBOHD mutant constructs, the coding regions of RBOHD (S8A, S39A, S611A, S703A, or S862A) were cloned into *Bam*HI site of pcDNA3.1-3FLAG-MCS [Kozak-3FLAG-*Bam*HI-*Eco*RV-stop fragment was inserted between *NheI* and *KpnI* sites of pcDNA3.1(-) vector (Invitrogen)]. Amino acid substituted mutants of CRK2 and RBOHD were generated by point-mutant primers and the mega-primer PCR method. pEF1-3Myc-GFP (Kawarazaki et al., 2013), pcDNA3.1-3FLAG-RBOHD, pcDNA3.1-3FLAG-RBOHC, pcDNA3.1-3FLAG-RBOHF and pcDNA3.1-3FLAG-MCS were described previously (Kaya et al., 2019). Primer sequences are listed in the Table S4.

### Subcellular protein localization

Fluorescent images were obtained using a Leica TCS SP5 II HCS confocal microscope. For investigation of CRK2-YFP localization, 514 nm excitation and 525-590 nm detection range were used.

### ROS measurements

Leaf discs were collected using a cork borer from 4-week-old Arabidopsis plants and floated overnight in sterile distilled water in 96 well plates under continuous light (17 μmol m^-2^ s^-1^, LED) at room temperature. On the following day, water was replaced with assay buffer containing 34 mg/L luminol sodium salt (Sigma, #A4685), 20 mg/L horse radish peroxidase (Fujifilm Wako, #169-10791), 200 nM flg22 (GenScript), 200 μg/mL Chitin (Sigma, #C9752) or 1 μM AtPep1 (ATKVKAKQRGKEKVSSGRPGQHN: synthesized by Synpeptide, China). Luminescence was measured for 1 sec every 1 min at room temperature using GloMax-Multi+Detection System (Promega). ROS production was expressed in relative luminescence units (RLU).

The ROS producing activity of RBOHs in HEK293T cells was measured as described previously (Kimura et al., 2012). Two days after transfection, medium was removed and cells were gently washed with 1×HBSS (GIBCO, #14025-092 or #14175-095). Measurements were started after addition of the assay buffer containing 250 μM lunimol sodium salt and 66.7 mg/L horse radish peroxidase. After 30 min measurement, 1 μM ionomycin (Calbiochem, #407952) was added. Chemiluminescence was measured for 1 sec every 1 min at 37 °C using GloMax-Multi+Detection System. ROS production was expressed in relative luminescence units (RLU). Expressed proteins were detected by immunoblotting with anti-FLAG (Sigma, #F1804), anti-cMyc (Fujifilm Wako, #017-2187), anti-β-actin (Sigma, #A5316) and IRDye800CW antimouse IgG (LI-COR, #926-32210) antibodies.

### Bacterial growth assay

To quantify bacterial growth on 4-week-old plants infected with the virulent *Pto*DC3000 (Whalen et al., 1991), growth curve assays were performed as described previously (Wrzaczek et al., 2007). Spray infections were carried out as previously described (Yao et al., 2013).

### Stomatal aperture

Measurements of stomatal aperture were carried out as previously described (Kadota et al., 2014) with minor modifications. Leaves were cut at the petiole from 3-week-old plants and incubated in water for 2 h. Samples were then incubated overnight in stomatal opening buffer (50 mM KCl, 10 μM CaCl_2_, 0.01% Tween20, 10 mM MES pH 6.15) to induce stomatal opening. For treatments, 5 μM flg22 was added to the buffer for 2 h prior to imaging. Brightfield images were obtained using a Leica TCS SP5 II HCS confocal microscope.

### Ca^2+^ imaging

Calcium imaging with YCNano-65 expressing plants was performed as described previously (Choi et al., 2014, Lenglet et al., 2017, Toyota et al., 2018). In brief, YCNano-65 was visualized by a fluorescence stereo microscope (Nikon) with a 1× objective lens (Nikon), image splitting optics (Hamamatsu Photonics) and a sCMOS camera (Hamamatsu Photonics). To excite YCNano-65, a mercury lamp (Nikon), a 436/20 nm excitation filter (Chroma) and a 455 nm dichroic mirror (Chroma) were used. The fluorescent signal from YCNano-65 was separated by a 515 nm dichroic mirror (Chroma) equipped in the image splitting optics. The resultant CFP and YFP (FRET) signals passed independently through a 480/40 nm and 535/30 nm emission filters, respectively (Chroma). A pair of the CFP and FRET images was simultaneously acquired every 4 s with the sCMOS camera using NIS-Elements imaging software (Nikon). A 7-day-old seedling was filled with approximately 50 μL of 1 μM flg22 or 1× MS 1 % sucrose liquid media. A region of interest (ROI) was placed on the adaxial surface of cotyledons to analyze both CFP and FRET signals. The FRET/CFP ratio was calculated by the 6D imaging plug-in modules in NIS-Elements imaging software (Nikon).

### MAPK assay

MAPK assays were performed as previously described (Yadeta et al., 2017). In brief, 4-week-old Arabidopsis plants were sprayed with 10 μM flg22 with 0.025 % Silwett L-77. Leaf samples were ground in liquid nitrogen and sand. Extraction buffer [50 mM HEPES (pH7.4), 50 mM NaCl, 10 mM EDTA, 0.2 % Triron X-100, 1 % Protease inhibitor cocktail (SIGMA, #P9599), 1 % Halt phosphatase inhibitor cocktail (Thermo scientific, #78428)] was added (2 mL/g plant powder). Samples were incubated at 4 °C for 30 min and centrifuged at 12,000 × g, 4 °C for 10 min. The supernatant was used for immunoblotting with anti-Phospho-p44/42 MAPK (Cell Signaling Technology, #4370) and IRDye800CW anti-rabbit IgG (LI-COR, #926-32211) antibodies.

### Callose Staining

Callose staining was performed as described previously (Hunter et al., 2019).

### qRT-PCR

Col-0, *crk2* and *fls2* seedlings were grown on MS 1 % sucrose agar plate for 5 days and were transferred into MS 1 % sucrose liquid media and grown for 5 days. Plants were incubated with 1 μM flg22 for 30 min, 1 h and 3 h, respectively. Plants were ground in liquid nitrogen and total RNA was extracted using the GeneJET Plant RNA purification Kit (Thermo scientific, #K0802). Total RNA was treated with DNase I (Thermo scientific, #EN0525) and cDNA was synthesized with Maxima H Minus Reverse Transcriptase (Thermo scientific, #EP0751). qPCR analysis was performed with CFX real-time PCR (BioRad, Hercules, CA, US) using 5× HOT FIREPol EvaGreen qPCR Mix Plus ROX (Soils Biodyne). *SAND, TIP41* and *YLS8* were used as reference genes for normalization. Relative expression was calculated with qBase+ (Biogazelle; https://www.qbaseplus.com/). Primers are listed in Table S4.

### Phytohormone analysis

Salicylic acid (SA) was analyzed from adult rosettes as described previously (Forcat et al., 2008) with minor modifications. Rosettes were flash-frozen in liquid nitrogen and freeze-dried for 24 h. Rosettes were ground in a mortar before weighing in to 2 mL tubes. About 6 mg aliquots of freeze-dried material were further homogenized by shaking with 5 mm stainless steel beads in a Qiagen Tissue Lyser II for 2 min at 25 Hz. Shaking was repeated after addition of 400 μL extraction buffer (10 % methanol, 1 % acetic acid) with internal standard (28 ng Salicylic-d4 Acid; CDN Isotopes, Pointe-Claire, Canada). Samples were then incubated on ice for 30 min and centrifuged for 10 min at 16,000 × g and 4 °C. Supernatants were centrifuged 3 times to remove all debris before LC-MS/MS analysis. The chromatographic separation was carried out using an Acquity UHPLC Thermo system(Waters, Milford, U.S.) equipped with a Waters Cortecs C18 column (2.7 μm, 2.1 × 100 mm). The solvent gradient (acetonitrile (ACN) / water with 0.1 % formic acid each) was adapted to a total run time of 7 min: 0-4 min 20 % to 95 % ACN, 4-5 min 95 % ACN, 5-7 min 95 % to 20 % ACN; flow rate 0.4 mL / min. For hormone identification and quantification, a tandem mass spectrometric system Xevo TQ-XS, triple quadrupole mass analyser (QqQ) with a ZSpray ESI function (Waters, Milford, U.S.) was used. Mass transitions were: SA 137 > 93, D4-SA 141 > 97.

### Protein extraction and Co-immunoprecipitation

Co-immunoprecipitation was performed as described previously (Kadota et al., 2016). Homozygous *35S::FLAG-RBOHD/rbohD* was crossed with homozygous *35S::CRK2-YFP/Col-0* or *35S::YFP-6Myc/Col-0. 35S::FLAG-RBOHD/35S::CRK2-YFP/rbohD* F3 plants were selected by kanamycin resistance (homozygous FLAG-RBOHD insertion) and PCR (homozygous *rbohD* T-DNA insertion). F1 and F3 plants were grown on MS 1 % sucrose agar plate for 7 days and were transferred into MS 1 % sucrose liquid media and grown for 8-10 days. F3 plants were incubated in water or 1 μM flg22 for 10 min or 30 min after vacuum application for 2 min. Plants were ground in liquid nitrogen and sand. Extraction buffer [50 mM Tris-HCl (pH 7.5), 150 mM NaCl, 10 % Glycerol, 5 mM DTT, 1 % Protease inhibitor cocktail (SIGMA, P9599), 2 % IGEPAL CA630, 1 mM Na_2_MoO_4_2H_2_O, 2.5 mM NaF, 1.5 mM Activated sodium orthovanadate, 1 mM PMSF] was added at 1.5 - 2 mL/g fresh weight. Samples were incubated at 4 °C for 1 h and centrifuged at 15,000 × g, 4 °C for 20 min. Supernatants were adjusted to 5 mg/mL protein concentration and incubated for 1 h at 4 °C with 100 μL of anti-GFP magnetic beads (Miltenyi Biotec, #130-091-125). Bound proteins were analyzed by immunoblotting with anti-GFP (Invitrogen, #A11122), anti-RBOHD (Agrisera, #AS15-2962), and IRDye800CW anti-rabbit IgG (LI-COR, #926-32211) antibodies.

To detect CRK2-YFP, total protein was extracted from 13-day-old *CRK2pro::CRK2(WT, K353E or D450N)/crk2* T3 homozygous plants with the same extraction buffer and analyzed by immunoblotting with anti-GFP (Invitrogen, #A11122) and IRDye800CW anti-rabbit IgG (LI-COR, #926-32211) antibodies. GFP signal intensity was quantified by ImageJ and normalized by CBB staining intensity.

To detect 3FLAG-RBOHD, total protein was extracted from 14-day-old *RBOHDpro::3FLAG-RBOHD (WT or S703A)/rbohD* T3 homozygous plants with the same extraction buffer and analyzed by immunoblotting with anti-FLAG (Sigma, #F1804 and IRDye800CW anti-mouse IgG (LI-COR, #926-32210) antibodies.

### Protein purification from *E.coli*

Cytosolic regions of CRK2 were expressed in *Escherichia coli* Lemo21. Cytosolic regions of RBOHD and Maltose binding protein (MBP) were expressed in *Escherichia coli* BL21. Glutathione S transferase (GST)-tagged recombinant proteins were purified using glutathione sepharose 4B (GE Healthcare, #17-0756-01) according to manufacturer’s instructions. MBP-tagged proteins were purified using amylose resin (New England Biolabs, #E8021S) according to manufacturer’s instructions.

### *In vitro* pull down

6His-GST-CRK2cyto, 6His-MBP-RBOHD/N, 6His-MBP-RBOHD/C and MBP were incubated with glutathione Sepharose 4B in the pull down buffer (20 mM HEPES, 50 mM KCl, 5 mM MgCl_2_, 1 % Tween20, 1 mM DTT and 100 μM PMSF) at 4 °C for 1 h. The glutathione sepharose 4B was washed four times with the pull down buffer and eluted with 10 mM reduced gluthatione. The mixture was analyzed by immunoblotting anti-6×His (Invitrogen, #MA1-135), anti-MBP (Santa Cruz Biotechnology, #sc-13564) and IRDy800CW anti-mouse IgG antibodies.

### *In vitro* kinase assay

Purified recombinant proteins were incubated with [γ-^32^P] ATP for 30 min at room temperature in the kinase assay buffer [50 mM HEPES (pH7.4), 1 mM DTT, 10 mM MgCl_2_, 0.6 mM unlabeled ATP]. The mixture was subsequently separated by SDS-PAGE and autoradiography was detected by FLA-5100 image analyzer (Fujifilm, Japan). For identification of *in vitro* phosphorylation sites by LC-ESI-MS/MS, 1.5 mM unlabeled ATP was used in the kinase buffer. The proteins were separated by SDS-PAGE, followed by CBB staining and were digested by trypsin (Thermo scientific, #90057) or Lys-C (Thermo scientific, #90051).

### Identification of *in vitro* phosphorylation sites of RBOHD by LC-ESI-MS/MS

Trypsin or Lys-C digested protein samples were analyzed by a Q Exactive mass spectrometer (Thermo Fisher Scientific, Bremen, Germany) connected to Easy NanoLC 1000 (Thermo Fisher Scientific). Peptides were first loaded on a trapping column and subsequently separated inline on a 15 cm C18 column (75 μm × 15 cm, ReproSil-Pur 5 μm 200 Å C18-AQ, Dr. Maisch HPLC GmbH, Ammerbuch-Entringen, Germany). The mobile phase consisted of water with 0.1 % formic acid (solvent A) or acetonitrile/water [80:20 (v/v)] with 0.1 % formic acid (solvent B). A linear 10 min gradient from 8 % to 42 % B was used to elute peptides.

MS data was acquired automatically by using Thermo Xcalibur 3.1 software (Thermo Fisher Scientific). An information dependent acquisition method consisted of an Orbitrap MS survey scan of mass range 300-2000 m/z followed by HCD fragmentation for 10 most intense peptide ions. Raw data was searched for protein identification by Proteome Discoverer (version 2.2) connected to in-house Mascot (v. 2.6.1) server.

Phosphorylation site locations were validated using phosphoRS algorithm. A SwissProt database with a taxonomy filter *Arabidopsis thaliana* was used. Two missed cleavages were allowed. Peptide mass tolerance ± 10 ppm and fragment mass tolerance ± 0.02 Da were used. Carbamidomethyl (C) was set as a fixed modification and methionine oxidation, acetylation of protein N-terminus, phosphorylation of Ser and Thr were included as variable modifications. Only peptides with FDR 0.01 were used.

### Targeted (phospho) peptide analysis

#### Plant treatment

Arabidopsis seeds were sterilized by incubating with 1.5 % NaClO 0.02 % Triton X-100 solution for 5 min and vernalized at 4 °C for 2 days. Sterilized seeds were germinated and grown in liquid culture on 6 well plates (30 seeds/well) in MGRL medium with 0.1 % (w/v) sucrose (2 mL/well) (Fujiwara et al., 1992) at 23 °C under continuous light (100 μmol m^-2^ s^-1^, LED) in a Percival growth chamber. Plates with 11-day-old seedlings were transferred from the growth chamber to a workbench and kept overnight for acclimatization before treatments. Seedlings were treated with either 1 μM flg22 or sterile water for 5 min after which seedlings were immediately collected and flash-frozen in liquid nitrogen and stored at −80 °C. Leaf discs were collected using a cork borer from 4-week-old Arabidopsis plants and floated overnight in sterile distilled water in petri dishes under continuous light (17 μmol m^-2^ s^-1^, LED) at room temperature. On the following day, water was replaced with new sterile distilled water or 200 nM flg22. After 5 min incubation leaf discs were immediately collected and flash-frozen in liquid nitrogen and stored at −80 °C.

#### Phosphopeptide enrichment

Frozen seedlings or leaf discs were disrupted using a Retsch mill (5 min, 30 Hz), and 500 μL urea extraction buffer [8 M urea in 100 mM Tris, pH 8.5, 20 μL/mL Phosphatase Inhibitor Cocktail 3 (Sigma, P0044), 20 μL/mL Phosphatase Inhibitor Cocktail 2 (Sigma, P5726), 5 mM DTT] was added to the disrupted frozen powders, mixed briefly and incubated at RT for 30 min. After centrifugation at 15,000 × g for 10 min, supernatants were transferred to fresh tubes. Protein concentrations were determined using Pierce 660 nm protein assay (Thermo Scientific). Extracts with 500 μg of protein were alkylated with 14 mM chloroacetamide (CAA) at RT for 30 min in the dark, CAA was quenched by addition of 1/200 sample volume 1M DTT. Samples were diluted 1:8 with 0.1 M Tris, pH 8.5, 1 mM CaCl_2_ and were digested overnight at RT either with 5 μg trypsin or 5 μg Lys-C. Digestion reaction was terminated by addition of TFA (0.1 % final concentration), and peptides were desalted using C18 SepPaks [1cc cartridge, 100 mg (WAT023590)]. In brief, SepPaks were conditioned using methanol (1 mL), buffer B (80 % acetonitrile, 0.1 % TFA; 1 mL) and buffer A (0.1 % TFA; 2 mL). Samples were loaded by gravity flow, washed with buffer A (1 × 1 mL, 1 × 2 mL) and eluted with buffer B (2 × 400 μL). 40 μL of eluates were kept separately to measure non-phosphopeptides and the rest were used for further phosphopeptide enrichment. Phosphopeptide enrichment was performed by hydroxy acid-modified metal-oxide chromatography (HAMMOC) using titania as described previously with minor modifications (Nakagami, 2014; Sugiyama et al., 2007).

#### LC-MS/MS data acquisition

Samples were analyzed using an EASY-nLC 1200 (Thermo Fisher) coupled to a Q Exactive Plus mass spectrometer (Thermo Fisher). Peptides were separated on 16 cm frit-less silica emitters (New Objective, 0.75 μm inner diameter), packed in-house with reversed-phase ReproSil-Pur C18 AQ 1.9 μm resin (Dr. Maisch). Peptides were loaded on the column and eluted for 115 min using a segmented linear gradient of 5 % to 95 % solvent B (0 min, 5 % B; 0-5 min, 5 % B; 5-65 min, 20 % B; 65-90 min, 35 % B; 90-100 min, 55 % B; 100-105 min, 95 % B; 105-115 min, 95 % B) [solvent A (0 % ACN, 0.1 % FA); solvent B (80 % ACN, 0.1 % FA)] at a flow rate of 300 nL/min. Mass spectra were acquired using a targeted (parallel reaction monitoring, PRM) approach. The acquisition method consisted of a full scan method combined with a non-scheduled PRM method. The 16 targeted precursor ions were selected based on the results of a DDA peptide search of phospho-enriched samples in Skyline (MacLean et al., 2010) (Version 4.2.0.x, https://skyline.ms). MS spectra were acquired in the Orbitrap analyzer with a mass range of 300–1750 m/z at a resolution of 70,000 FWHM and a target value of 3×10^6^ ions, followed by MS/MS acquisition for the 16 targeted precursors. Precursors were selected with an isolation window of 2.0 m/z. HCD fragmentation was performed at a normalized collision energy of 27. MS/MS spectra were acquired with a target value of 2×10^5^ ions at a resolution of 17,500 FWHM, a maximum injection time of 120 ms and a fixed first mass of m/z 100.

#### MS data analysis

Raw data from PRM acquisition were processed using MaxQuant software (version 1.5.7.4, http://www.maxquant.org/) (Cox and Mann, 2008). MS/MS spectra were searched by the Andromeda search engine against a combined database containing the sequences from *Arabidopsis thaliana* (TAIR10_pep_20101214; ftp://ftp.arabidopsis.org/home/tair/Proteins/TAIR10_protein_lists/) and sequences of 248 common contaminant proteins and decoy sequences. Trypsin specificity was required and a maximum of two missed cleavages allowed. Minimal peptide length was set to seven amino acids. Carbamidomethylation of cysteine residues was set as fixed, phosphorylation of serine, threonine and tyrosine, oxidation of methionine and protein N-terminal acetylation as variable modifications. The match between runs option was disabled. Peptide-spectrum-matches and proteins were retained if they were below a false discovery rate of 1 % in both cases. The “msms.txt” output from MaxQuant was further analyzed using Skyline in PRM mode. Trypsin specificity was required and a maximum of two missed cleavages allowed. Minimal and maximum peptide lengths were set to 7 and 25 amino acids, respectively. Carbamidomethylation of cysteine, phosphorylation of serine, threonine and tyrosine, oxidation of methionine, and protein N-terminal acetylation were set as modifications. Results were filtered for precursor charges of 2 and 3, and b- and y-ions with ion charges of +1 and +2. Product ions were set to “from ion 1 to last ion”. All chromatograms were inspected manually and peak integration was corrected for best representation of MS2 signals. Peak area data was exported and further processed. The Skyline documents containing the data for the targeted phophoproteomics experiments have been uploaded to Panorama Public and can be obtained from https://panoramaweb.org/RBOHDphosphorylation.url. Raw data have been deposited to the ProteomeXchange Consortium via the Panorama partner repository with the dataset identifier PXD013525 (http://proteomecentral.proteomexchange.org/cgi/GetDataset?ID=PXD013525).

### Phylogenetic analysis

Sequences for plant *RBOH* genes were extracted from public genome databases and manually curated. Sequences were aligned using MAFFT version 7.407 (Katoh and Standley, 2013) using iterative refinement (L-INS-i). This alignment was used to infer a phylogenetic maximum likelihood tree using RAxML version 8.1.3 (Stamatakis, 2014) with the JTT or DAYHOFF substitution models (which resulted in identical phylogenetic trees). Thousand bootstrap replicates were calculated using RAxML. Human NOX2 and NOX5ß were used as outgroup to root the phylogenetic tree of plant NADPH oxidases. The sequence alignment of plant RBOHs, human NOX2 and NOX5ß can be viewed on the (Veidenberg et al., 2016) webserver (http://was.bi?id=p_nwi1) and the tree as nexus file is available in Supplementary data. Sequence motifs were analyzed using the MEME suite (Bailey et al., 2009).

### Statistical analysis

Statistical analyses were performed with JMP Pro13 (SAS, https://www.jmp.com/). One-way ANOVA results are in Table S5.

## Data availability

Phylogenetic tree of human and plant NADPH oxidases with bootstrap information for 1000 replicates and corresponding sequence alignment has been deposited on Wasabi (http://was.bi?id=pnwi1). Data for the targeted phophoproteomics experiments has been uploaded to Panorama Public (https://panoramaweb.org/RBOHDphosphorylation.url). Raw data have been deposited to the ProteomeXchange Consortium via the Panorama partner repository with the dataset identifier PXD013525 (http://proteomecentral.proteomexchange.org/cgi/GetDataset?ID=PXD013525). Materials used in the experimental work are available from the authors upon request.

## Author contributions

SK, KH, HN and MW conceived and designed the project. SK, KH, LV, CT, MC, AV, AR, LM, MM, TW, MT, and MW carried out experiments. SK, KH, LV, AV, AR, TH, MT, and MW analyzed the data. AH, SCS and HN designed and performed targeted MS analysis and analyzed the data. SK and MW wrote the manuscript. All authors read and contributed to the final manuscript.

## Acknowledgments

The authors would like to thank Dr. Julia Krasensky-Wrzaczek and Dr. Alexey Shapiguzov (University of Helsinki, Finland) for critical comments on the manuscript. We thank Tuomas Puukko, Nghia Le Tri, Simon Schmitz, Denis Owczarek, Jan-Niklas Weber (University of Helsinki, Finland) and Jiaqi Wang (Saitama University, Japan) for technical assistance, Dr. Riccardo Siligato for the Gateway Multisite vector system. LV and TH thank Trude Johansen for technical assistance in LC-MS/MS based hormone analysis. We thank Dr. Yasuhiro Kadota (RIKEN Yokohama, Japan) and Prof. Cyril Zipfel (University of Zurich, Switzerland) for *35S::FLAG-RBOHD/rbohD* seeds and pOPINM-RBOHD/N and pBin19g-pRBOHD::3FLAG-RBOHD plasmids. We thank Prof. Simon Gilroy (University of Wisconsin, USA) for *YCNano65* seeds. The pcDNA3.1-3FLAG-RBOHD and pcDNA3.1-3FLAG-MCS plasmids were provided by Prof. Kazuyuki Kuchitsu (Tokyo University of Science, Japan). Microscopy imaging was performed at the Light Microscopy Unit, Institute of Biotechnology, University of Helsinki. Mass spectrometry analyses were performed at the Turku Proteomics Facility, supported by Biocenter Finland. This work was supported by the Academy of Finland (grant numbers #275632, #283139, #312498, and #323917 to MW), the University of Helsinki (Three-year fund allocation to MW), the Finnish Cultural Foundation (grant numbers 00170046 and 00181379 to LV), and KAKENHI (17H05007, 18H04775, and 18H05491 to MT), the Max-Planck-Gesellschaft (to HN). KH, SK, AV and MW are members of the Centre of Excellence in the Molecular Biology of Primary Producers (2014-2019) funded by the Academy of Finland (grant numbers #271832 and #307335).

The authors declare that they have no conflict of interest.

## Supplemental figure legends

**Supplemental Figure 1. Complementation of *crk2* with *CRK2pro::CRK2-YFP* (Supports Figure 1)**.

**(A)** Box plot shows dry weight of 21-day-old plants (n = 10). Differences compared with Col-0 were evaluated with One-way Anova with Tukey-Kramer HSD, *** p < 0.001, ns, not statistically significant. The experiment was repeated three times with similar results.

**(B)** Box plot shows salicylic acid accumulation level in Col-0, *crk2, CRK2pro::CRK2-YFP/crk2 #1-22* and C24 (n ≥ 19). 25-day-old plants were used. The experiment was repeated five times (three or four data points per experiment) and all data points are blotted. Differences compared with Col-0 were evaluated with Oneway Anova with Tukey-Kramer HSD, ** p < 0.01, ns, not statistically significant.

**(C)** and **(D)** Subcellular localization of CRK2-YFP, CRK2^K353E^-YFP and CRK2^D450N^-YFP of 7-day-old seedlings.

**(C)** In leaves. Plasma membrane localization was confirmed using plasmolysis to visualize Hechtian strands (arrow head). Plasmolysis was induced by the application of 0.8 M mannitol. Scale bar = 25 μm.

**(D)** In roots. Scale bar = 20 μm.

**(E)** Expression of CRK2-YFP, CRK2^K353E^-YFP and CRK2^D450N^-YFP in 13-day-old seedlings were detected by anti-GFP antibody. Input proteins stained with amido black staining (lower panel). To quantify the expression level, GFP signal intensity was normalized by amido black signal intensity (GFP/amido black value).

**Supplemental Figure 2. MAMP-triggered ROS production and molecular responses in *crk2* (Supports Figure 2)**.

**(A)** and **(B)** Box plot shows quantitative real-time RT-PCR (qPCR) analysis of *FRK1* **(A)** or *NHL10* **(B)** transcripts in Col-0, *crk2* and *fls2* after treatment with flg22 (n = 3, biological replicates). 10-day-old plants were incubated in 1 μM flg22 solution and collected at indicated time (each data point contains 90 plants). Sample collection and qPCR were repeated three times with the same procedure and the three independent results were plotted. Transcript levels were calculated by comparison with non-treated Col-0 (Time = 0). Different letters indicate significant difference at p < 0.05 (One-way Anova with Tukey-Kramer HSD).

**(C)** and **(D)** Chitin- or AtPep1-induced ROS production in Col-0, *crk2* and *rbohD.* Leaf discs from 28-day-old plants were treated with 200 μg/mL chitin (**C**) or 1 μM AtPep1 (**D**). ROS production is expressed in relative luminescence units (RLU). Box plots show integration of ROS production for 40 min (upper right). Differences compared with Col-0 were evaluated with One-way Anova with Tukey-Kramer HSD. The experiment was repeated three times with similar results. **(C)** Values represent the mean ±SEM of n ≥ 21, * p < 0.05, *** p < 0.001. **(D)** Values represent the mean ±SEM of n = 24, *** p < 0.001.

**(E)** Quantification of flg22-induced stomatal closure in Col-0 (n = 28) and *crk2* (n ≥ 40). Leaf discs from 21-day-old plants were treated with 5 μM flg22 for 2 h. Letters indicate significant differences at p < 0.05 (Oneway Anova with Tukey-Kramer HSD).

**(F)** Quantitative analysis of cytosolic Ca^2+^ changes in response to 1×MS 1 % sucrose liquid media or 1 μM flg22 in 7-day-old *YCNano65* or *YCNano65/crk2* seedlings. Values represent the mean ±SEM of n = 9 *(YCNano65)* or n = 15 *(YCNano65/crk2).*

**(G)** Representative frame images of cytosolic Ca^2+^ change in wild type and *crk2* plants. Bar = 0.5 mm.

**(H)** MAPK activation in Col-0, *crk2* and *fls2* in response to treatment with 1 μM flg22. 28-day-old plants (12 plants per genotype). Phosphorylated MPK3 and MPK6 were detected with anti-p44/42 MPK antibody (upper panel). Proteins stained with amido black staining (lower panel). The experiment was repeated three times with similar results.

**(I)** Quantification of flg22-induced callose deposition by aniline blue (n ≥ 16) in 7-day-old seedlings without (nt), 30 minutes (30min) or 12 hours (12h) treatment with 10 μM flg22. Letters indicate significant differences at p < 0.05 (One-way Anova with Tukey-Kramer HSD).

**(J)** Representative images of aniline blue stained leaves. Bar = 100 μm.

**Supplemental Figure 3. CRK2 modulates the ROS-producing activity of RBOHC, D and F in HEK293T cells (Supports Figure 3).**

**(A)** Expressed proteins were detected by anti-FLAG and anti-Myc antibodies (Figure 3A). 3FLAG-RBOHD: 107 kDa, CRK2-3Myc: 75.8 kDa, 3Myc-GFP: 31 kDa, β-actin 42 kDa. As a loading control, β-actin was used. Loading volume for anti-Myc antibody: 3FLAG-RBOHD + 3Myc-GFP (5 μL), the others (50 μL).

**(B)** ROS production in RBOHD-expressing HEK293T cells in Ca^2+^-free buffer. 3FLAG-RBOHD was transiently co-expressed with either 3Myc-GFP or 3Myc-CRK2 (WT or D450N) in HEK293T cells. Values represent mean ±SEM of n = 3. E.V. = empty vector. The experiment was repeated three times with similar results.

**(C)** Proteins expressed in HEK293T cells were detected by anti-FLAG and anti-Myc antibodies (Supplemental Figure 3B). 3FLAG-RBOHD: 107 kDa, CRK2-3Myc: 75.8 kDa, 3Myc-GFP: 31 kDa, β-actin 42 kDa. As a loading control, β-actin was used. Loading volume for anti-Myc antibody: 3FLAG-RBOHD + 3Myc-GFP (5 μL), others (50 μL).

**(D)** to **(F)** ROS production of RBOHD-, RBOHC-, or RBOHF-expressing HEK293T cells. 3FLAG-RBOHD

**(D)**, 3FLAG-RBOHC **(E)**, or 3FLAG-RBOHF **(F)** was transiently co-expressed with 3Myc-GFP or CRK2-3Myc in HEK293T cells, respectively. After 20 min of base line measurement, 1 μM ionomycin was added to the medium (black arrow). Values represent mean ±SEM of n = 3. The experiment was repeated two times with similar results.

**(G)** Proteins expressed in HEK293T cells were detected by anti-FLAG and anti-Myc antibodies (Supplemental Figures 3D to 3F). 3FLAG-RBOHD: 107 kDa, 3FLAG-RBOHC, 106 kDa, 3FLAG-RBOHF: 111 kDa, CRK2-3Myc: 75.8 kDa, 3Myc-GFP: 31 kDa, β-actin 42 kDa. As a loading control, β-actin was used. Loading volume for anti-Myc antibody: 3FLAG-RBOHs + 3Myc-GFP (5 μL), 3FLAG-RBOHs + CRK2-3Myc (50 μL).

**Supplemental Figure 4. ROS production activity of RBOHD S703A and S862A in HEK293T cells (Supports Figure 5).**

**(A)** and **(B)** Proteins expressed in HEK293T cells were detected by anti-FLAG and anti-Myc antibodies (Figures 5A and 5B). 3FLAG-RBOHD: 107 kDa, CRK2-3Myc: 75.8 kDa, 3Myc-GFP: 31 kDa, β-actin 42 kDa. As a loading control, β-actin was used. Loading volume for anti-Myc antibody: 3FLAG-RBOHD + 3Myc-GFP (5 μL), others (50 μL).

**(A)** 3FLAG-RBOHD (WT, S8A or S39A) was transiently co-expressed with either 3Myc-GFP or CRK2-3Myc into HEK293T cells (Figure 5A).

**(B)** 3FLAG-RBOHD (WT, S611A, S703A or S862A) was transiently co-expressed with either 3Myc-GFP or CRK2-3Myc into HEK293T cells (Figure 5B).

**(C)** and **(D)** ROS production of RBOHD-expressing HEK293T cells. After 30 min 1 μM ionomycin was added to the medium (black arrow). Values represent mean ±SEM of n = 3. E.V. = empty vector. The experiment was repeated three times with similar results.

**(C)** 3FLAG-RBOHD (WT or S703A) was transiently co-expressed with either 3Myc-GFP or CRK2-3Myc into HEK293T cells. The right upper panel is the enlargement of ionomycin-induced transient ROS production of 3FLAG-RBOHD (WT or S703A) and 3Myc-GFP co-expressing cells (dashed box).

**(D)** 3FLAG-RBOHD (WT or S862A) was transiently co-expressed with either 3Myc-GFP or CRK2-3Myc into HEK293T cells.

**Supplemental Figure 5. Quantification of RBOHD and MPK phosphorylation upon flg22 treatment (Supports Figure 6).**

**(A)** to **(J)** 12-day-old Col-0 seedlings were treated with water (-) or 1 μM flg22 (+) for 5 min. Total proteins were digested by trypsin for peptides from RBOHD N-terminal region and MPKs, by Lys-C for RBOHD S703 peptide. Peptides were enriched, and then selected phosphopeptides were quantified by LC-MS/MS. Box plots show MS2 fragment peak ion areas of indicated phosphopeptides (n = 4). Differences between water- or flg22-treated samples were evaluated with One-way Anova with Tukey-Kramer HSD, * P<0.05, ** P<0.01, *** p<0.001, ns, not statistically significant.

**(A)** and **(B)** RBOHD S8 residue.

**(C)** RBOHD S39 residue.

**(D)** RBOHD S703 residue.

**(E)** RBOHD S163 residue.

**(F)** RBOHD S347 residue.

**(G)** RBOHD S343 and S347 residues.

**(H)** MPK3 TEY motif.

**(I)** MPK6 TEY motif.

**(J)** MPK11 TEY motif.

**(K)** to **(M)** Quantification of RBOHD S8 and S39 phosphorylation in Col-0, *CRK2pro::CRK2-YFP/crk2* #1-22 and *CRK2pro::CRKř^450N^-YFP/crk2* #7-2 upon flg22 treatment. Leaf discs from 28-day-old plants were treated with water (-) or 200 nM flg22 (+) for 5 min. Total proteins were digested with trypsin and phosphopeptides were enriched, and then selected phosphopeptides were quantified by LC-MS/MS. Box plots show MS2 fragment peak ion areas of indicated phosphopeptides. Each data point contains 14 leaf discs. Letters indicate significant differences at p < 0.05 (One-way Anova with Tukey-Kramer HSD).

**(K)** RBOHD S8 residue (n ≥ 2).

**(L**) and (**M**) RBOHD S39 residue. n ≥ 2 (**L**), n = 3 (**M**).

**Supplemental Figure 6. RBOHD S703 is involved in regulation of flg22-induced ROS production (Supports Figure 6).**

**(A)** Representative pictures of 21-day-old plants of *RBOHDpro::3FLAG-RBOHD/rbohD* #1-3, #11-1 and *RBOHDpro::3FLAG-RBOHD^S703A^/rbohD* #3-2, #1-4 plants. Bar = 1 cm.

**(B)** and **(C)** Expressed proteins in 14-day-old *RBOHDpro::3FLAG-RBOHD/rbohD* and *RBOHDpro::3FLAG-RBOHD^S703Λ^/rbohD.* 3FLAG-RBOHD was detected by anti-FLAG antibody. Input proteins stained with amido black staining (lower panel).

**(B)** *RBOHDpro::3FLAG-RBOHD/rbohD* #1-3 and *RBOHDpro::3FLAG-RBOHD/rbohD* #3-2.

**(C)** *RBOHDpro::3FLAG-RBOHD/rbohD* #11-1 and *RBOHDpro::3FLAG-RBOHD^S703A^/rbohD* #1-4.

**(D)** flg22-induced ROS production in *RBOHDpro::3FLAG-RBOHD/rbohD* #11-1 and *RBOHDpro::3FLAG-RBOHD^S703A^/rbohD* #1-4. Leaf discs from 28-day-old plants were treated with 200 nM flg22. Box plots show integration of ROS production for 40 min (upper right). Values represent the mean ±SEM of n = 24. Difference between lines was evaluated with One-way Anova with Tukey-Kramer HSD, *** p < 0.001.

**(E)** Quantification of flg22-induced stomatal closure in *rbohD* (n ≥ 21)*, RBOHDpro::3FLAG-RBOHD/rbohD* #1-3 (n ≥ 35) and *RBOHDpro::3FLAG-RBOHD^S703A^/rbohD* #3-2 (n ≥ 35). Leaf discs from 21-day old plants were treated with 5 μM flg22 for 2 h. Letters indicate significant differences at p < 0.05 (One-way Anova with Tukey-Kramer HSD).

**Supplemental Figure 7. Reduced ROS production in *crk2* is not due to lower expression of *BIK1* (Supports discussion).**

Box plot shows quantitative real-time RT-PCR (qPCR) analysis of *BIK1* transcripts in Col-0, *crk2* and *fls2* after treatment with flg22 (n = 3, biological replicates). 10-day-old plants were incubated in 1 μM flg22 solution and collected at indicated time (each data point contains 90 plants). Sample collection and qPCR were repeated three times with the same procedure and the three independent results were plotted. Transcript levels were calculated by comparison with non-treated Col-0 (Time = 0). Different letters indicate significant difference at p < 0.05 (One-way Anova with Tukey-Kramer HSD).

## Supplemental table legends

**Supplemental Table 1. *In vitro* phosphorylation sites of 6His-MBP-RBOHDcyto by 6His-GST-CRK2cyto**

The 6His-MBP-RBOHD cytosolic regions were incubated with 6His-GST-CRK2cyto. The 6His-MBP-RBOHDcyto bands were excised from a SDS polyacrylamide gel and subsequently digested by trypsin or Lys-C. The peptides were analyzed by LC-MS/MS. Phosphorylated peptides are designated as pS.

**Supplemental Table 2. The numbers of Serine (S) and Threonine (T) residues in C-terminus of NADPH oxidases in plants and animals.**

**Supplemental Table 3. Progeny of *CRK2/crk2 BIK1/bik1* parent and *CRK2/crk2 bikl/bikl* parent**

The genotypes of F2 and F3 progenies were determined by PCR. Observed, the number of individuals observed; Expected, the expected number based on Mendelian inheritance. Chi-square test was used to determine the probability (P) of which the deviation of the observed value from the expected value was due to chance.

**Supplemental Table 4. Primer sequences**

**Supplemental Table 5. One-way ANOVA results**

## References

Acharya, B.R., Raina, S., Maqbool, S.B., Jagadeeswaran, G., Mosher, S.L., Appel, H.M., Schultz, J.C., Klessig, D.F., and Raina, R. (2007). Overexpression of CRK13, an Arabidopsis cysteine-rich receptor-like kinase, results in enhanced resistance to *Pseudomonas syringae*. Plant J 50, 488–499.

Bailey, T.L., Boden, M., Buske, F.A., Frith, M., Grant, C.E., Clementi, L., Ren, J., Li, W.W., and Noble, W.S. (2009). MEME SUITE: tools for motif discovery and searching. Nucleic Acids Res 37, W202–208.

Banfi, B., Tirone, F., Durussel, I., Knisz, J., Moskwa, P., Molnar, G.Z., Krause, K.H., and Cox, J.A. (2004). Mechanism of Ca^2+^ activation of the NADPH oxidase 5 (NOX5). J Biol Chem 279, 18583–18591.

Beaumel, S., Picciocchi, A., Debeurme, F., Vives, C., Hesse, A.M., Ferro, M., Grunwald, D., Stieglitz, H., Thepchatri, P., Smith, S.M.E., et al. (2017). Down-regulation of NOX2 activity in phagocytes mediated by ATM-kinase dependent phosphorylation. Free Radic Biol Med 113, 1–15.

Benschop, J.J., Mohammed, S., O’Flaherty, M., Heck, A.J., Slijper, M., and Menke, F.L. (2007). Quantitative phosphoproteomics of early elicitor signaling in *Arabidopsis*. Mol Cell Proteomics 6, 1198–1214.

Bigeard, J., Colcombet, J., and Hirt, H. (2015). Signaling mechanisms in pattern-triggered immunity (PTI). Mol Plant 8, 521–539.

Boudsocq, M., Danquah, A., de Zelicourt, A., Hirt, H., and Colcombet, J. (2015). Plant MAPK cascades: Just rapid signaling modules? Plant Signal Behav 10, e1062197.

Bourdais, G., Burdiak, P., Gauthier, A., Nitsch, L., Salojärvi, J., Rayapuram, C., Idänheimo, N., Hunter, K., Kimura, S., Merilo, E., et al. (2015). Large-scale phenomics identifies primary and fine-tuning roles for CRKs in responses related to oxidative stress. PLOS Genetics 11, e1005373.

Caillaud, M.C., Wirthmueller, L., Sklenar, J., Findlay, K., Piquerez, S.J., Jones, A.M., Robatzek, S., Jones, J.D., and Faulkner, C. (2014). The plasmodesmal protein PDLP1 localises to haustoria-associated membranes during downy mildew infection and regulates callose deposition. PLoS Pathog 10, e1004496.

Chen, D., Cao, Y., Li, H., Kim, D., Ahsan, N., Thelen, J., and Stacey, G. (2017). Extracellular ATP elicits DORN1-mediated RBOHD phosphorylation to regulate stomatal aperture. Nat Commun 8, 2265.

Chen, F., Yu, Y., Haigh, S., Johnson, J., Lucas, R., Stepp, D.W., and Fulton, D.J. (2014). Regulation of NADPH oxidase 5 by protein kinase C isoforms. PLoS One 9, e88405.

Chen, K., Fan, B., Du, L., and Chen, Z. (2004). Activation of hypersensitive cell death by pathogen-induced receptor-like protein kinases from *Arabidopsis*. Plant Mol Biol 56, 271–283.

Chern, M., Xu, Q., Bart, R.S., Bai, W., Ruan, D., Sze-To, W.H., Canlas, P.E., Jain, R., Chen, X., and Ronald, P.C. (2016). A genetic screen identifies a requirement for cysteine-rich-receptor-like kinases in rice NH1 (OsNPR1)-mediated immunity. PLoS Genet 12, e1006049.

Choi, W.G., Toyota, M., Kim, S.H., Hilleary, R., and Gilroy, S. (2014). Salt stress-induced Ca^2+^ waves are associated with rapid, long-distance root-to-shoot signaling in plants. Proc Natl Acad Sci USA 111, 6497–6502.

Clough, S.J., and Bent, A.F. (1998). Floral dip: a simplified method for *Agrobacterium-mediated* transformation of *Arabidopsis thaliana*. Plant J 16, 735–743.

Couto, D., and Zipfel, C. (2016). Regulation of pattern recognition receptor signalling in plants. Nat Rev Immunol 16, 537–552.

Cox, J., and Mann, M. (2008). MaxQuant enables high peptide identification rates, individualized p.p.b.-range mass accuracies and proteome-wide protein quantification. Nat Biotechnol 26, 1367–1372.

Dubiella, U., Seybold, H., Durian, G., Komander, E., Lassig, R., Witte, C.P., Schulze, W.X., and Romeis, T. (2013). Calcium-dependent protein kinase/NADPH oxidase activation circuit is required for rapid defense signal propagation. Proc Natl Acad Sci USA 110, 8744–8749.

Ellinger, D., Sode, B., Falter, C., and Voigt, C.A. (2014). Resistance of callose synthase activity to free fatty acid inhibition as an indicator of Fusarium head blight resistance in wheat. Plant Signal Behav 9, e28982.

Forcat, S., Bennett, M.H., Mansfield, J.W., and Grant, M.R. (2008). A rapid and robust method for simultaneously measuring changes in the phytohormones ABA, JA and SA in plants following biotic and abiotic stress. Plant Methods 4, 16.

Fujiwara, T., Hirai, M.Y., Chino, M., Komeda, Y., and Naito, S. (1992). Effects of sulfur nutrition on expression of the soybean seed storage protein genes in transgenic petunia. Plant Physiol 99, 263–268.

Han, J.P., Koster, P., Drerup, M.M., Scholz, M., Li, S., Edel, K.H., Hashimoto, K., Kuchitsu, K., Hippler, M., and Kudla, J. (2019). Fine-tuning of RBOHF activity is achieved by differential phosphorylation and Ca^2+^ binding. New Phytol 221, 1935–1949.

Hunter, K., Kimura, S., Rokka, A., Tran, H.C., Toyota, M., Kukkonen, J.P., and Wrzaczek, M. (2019). CRK2 enhances salt tolerance by regulating callose deposition in connection with PLDα1. Plant Physiol 180, 2004–2021.

Idänheimo, N., Gauthier, A., Salojärvi, J., Siligato, R., Brosché, M., Kollist, H., Mähönen, A.P., Kangasjärvi, J., and Wrzaczek, M. (2014). The *Arabidopsis thaliana* cysteine-rich receptor-like kinases CRK6 and CRK7 protect against apoplastic oxidative stress. Biochem Biophys Res Commun 445, 457–462.

Jagnandan, D., Church, J.E., Banfi, B., Stuehr, D.J., Marrero, M.B., and Fulton, D.J. (2007). Novel mechanism of activation of NADPH oxidase 5. Calcium sensitization via phosphorylation. J Biol Chem 282, 6494–6507.

Jiménez-Quesada, M.J., Traverso, J.A., and Alché Jde, D. (2016). NADPH oxidase-dependent superoxide production in plant reproductive tissues. Front Plant Sci 7, 359.

Kadota, Y., Macho, A.P., and Zipfel, C. (2016). Immunoprecipitation of plasma membrane receptor-like kinases for identification of phosphorylation sites and associated proteins. Methods Mol Biol 1363, 133–144.

Kadota, Y., Shirasu, K., and Zipfel, C. (2015). Regulation of the NADPH oxidase RBOHD during plant immunity. Plant Cell Physiol 56, 1472–1480.

Kadota, Y., Sklenar, J., Derbyshire, P., Stransfeld, L., Asai, S., Ntoukakis, V., Jones, J.D., Shirasu, K., Menke, F., Jones, A., et al. (2014). Direct regulation of the NADPH oxidase RBOHD by the PRR-associated kinase BIK1 during plant immunity. Mol Cell 54, 43–55.

Katoh, K., and Standley, D.M. (2013). MAFFT multiple sequence alignment software version 7: improvements in performance and usability. Mol Biol Evol 30, 772–780.

Kawarazaki, T., Kimura, S., Iizuka, A., Hanamata, S., Nibori, H., Michikawa, M., Imai, A., Abe, M., Kaya, H., and Kuchitsu, K. (2013). A low temperature-inducible protein AtSRC2 enhances the ROS-producing activity of NADPH oxidase AtRbohF. Biochim Biophys Acta 1833, 2775–2780.

Kaya, H., Takeda, S., Kobayashi, M.J., Kimura, S., Iizuka, A., Imai, A., Hishinuma, H., Kawarazaki, T., Mori, K., Yamamoto, Y., et al. (2019). Comparative analysis of the reactive oxygen species-producing enzymatic activity of Arabidopsis NADPH oxidases. Plant J 98, 291–300.

Kimura, S., Kaya, H., Kawarazaki, T., Hiraoka, G., Senzaki, E., Michikawa, M., and Kuchitsu, K. (2012). Protein phosphorylation is a prerequisite for the Ca^2+^-dependent activation of *Arabidopsis* NADPH oxidases and may function as a trigger for the positive feedback regulation of Ca^2+^ and reactive oxygen species. Biochim Biophys Acta 1823, 398–405.

Kimura, S., Waszczak, C., Hunter, K., and Wrzaczek, M. (2017). Bound by fate: The role of reactive oxygen species in receptor-like kinase signaling. Plant Cell 29, 638–654.

Lee, D.S.K., Young Cheon; Kwon, Sun Jae; Ryu, Choong-Min; Park, Ohkmae K. (2017). The Arabidopsis cysteine-rich receptor-like kinase CRK36 regulates immnity through interaction with the cytoplasmic kinase BIK1. Frontiers in Plant Science 8, 1856.

Lee, Y., Rubio, M.C., Alassimone, J., and Geldner, N. (2013). A mechanism for localized lignin deposition in the endodermis. Cell 153, 402–412.

Lee, Y., Yoon, T.H., Lee, J., Jeon, S.Y., Lee, J.H., Lee, M.K., Chen, H., Yun, J., Oh, S.Y., Wen, X., et al. (2018). A lignin molecular brace controls precision processing of cell walls critical for surface integrity in *Arabidopsis*. Cell 173, 1468–1480.

Lenglet, A., Jaslan, D., Toyota, M., Mueller, M., Müller, T., Schönknecht, G., Marten, I., Gilroy, S., Hedrich, R., and Farmer, E.E. (2017). Control of basal jasmonate signalling and defence through modulation of intracellular cation flux capacity. New Phytol 216, 1161–1169.

Li, L., Li, M., Yu, L., Zhou, Z., Liang, X., Liu, Z., Cai, G., Gao, L., Zhang, X., Wang, Y., et al. (2014). The FLS2-associated kinase BIK1 directly phosphorylates the NADPH oxidase RbohD to control plant immunity. Cell Host Microbe 15, 329–338.

Lin, Z.J., Liebrand, T.W., Yadeta, K.A., and Coaker, G. (2015). PBL13 Is a serine/threonine protein kinase that negatively regulates Arabidopsis immune responses. Plant Physiol 169, 2950–2962.

MacLean, B., Tomazela, D.M., Shulman, N., Chambers, M., Finney, G.L., Frewen, B., Kern, R., Tabb, D.L., Liebler, D.C., and MacCoss, M.J. (2010). Skyline: an open source document editor for creating and analyzing targeted proteomics experiments. Bioinformatics 26, 966–968.

Meitzler, J.L., Antony, S., Wu, Y., Juhasz, A., Liu, H., Jiang, G., Lu, J., Roy, K., and Doroshow, J.H. (2014). NADPH oxidases: a perspective on reactive oxygen species production in tumor biology. Antioxid Redox Signal 20, 2873–2889.

Miller, C.J., and Turk, B.E. (2016). Rapid identification of protein kinase phosphorylation site motifs using combinatorial peptide libraries. Methods Mol Biol 1360, 203–216.

Nakagami, H. (2014). StageTip-based HAMMOC, an efficient and inexpensive phosphopeptide enrichment method for plant shotgun phosphoproteomics. Methods Mol Biol 1072, 595–607.

Ogasawara, Y., Kaya, H., Hiraoka, G., Yumoto, F., Kimura, S., Kadota, Y., Hishinuma, H., Senzaki, E., Yamagoe, S., Nagata, K., et al. (2008). Synergistic activation of the *Arabidopsis* NADPH oxidase AtrbohD by Ca^2+^ and phosphorylation. J Biol Chem 283, 8885–8892.

Pandey, D., Gratton, J.P., Rafikov, R., Black, S.M., and Fulton, D.J. (2011). Calcium/calmodulin-dependent kinase II mediates the phosphorylation and activation of NADPH oxidase 5. Mol Pharmacol 80, 407–415.

Raad, H., Paclet, M.H., Boussetta, T., Kroviarski, Y., Morel, F., Quinn, M.T., Gougerot-Pocidalo, M.A., Dang, P.M., and El-Benna, J. (2009). Regulation of the phagocyte NADPH oxidase activity: phosphorylation of gp91^phox^/NOX2 by protein kinase C enhances its diaphorase activity and binding to Rac2, p67^phox^, and p47^phox^. FASEB J 23, 1011–1022.

Schmidt, R., Kunkowska, A.B., and Schippers, J.H. (2016). Role of reactive oxygen species during cell expansion in leaves. Plant Physiol 172, 2098–2106.

Shiu, S.H., and Bleecker, A.B. (2001). Receptor-like kinases from *Arabidopsis* form a monophyletic gene family related to animal receptor kinases. Proc Natl Acad Sci USA 98, 10763–10768.

Siligato, R., Wang, X., Yadav, S.R., Lehesranta, S., Ma, G., Ursache, R., Sevilem, I., Zhang, J., Gorte, M., Prasad, K., et al. (2016). MultiSite gateway-compatible cell type-specific gene-inducible system for plants. Plant Physiol 170, 627–641.

Stamatakis, A. (2014). RAxML version 8: a tool for phylogenetic analysis and post-analysis of large phylogenies. Bioinformatics 30, 1312–1313.

Stone, J.M., and Walker, J.C. (1995). Plant protein kinase families and signal transduction. Plant Physiol 108, 451–457.

Sugiyama, N., Masuda, T., Shinoda, K., Nakamura, A., Tomita, M., and Ishihama, Y. (2007). Phosphopeptide enrichment by aliphatic hydroxy acid-modified metal oxide chromatography for nano-LC-MS/MS in proteomics applications. Mol Cell Proteomics 6, 1103–1109.

Suzuki, N., Miller, G., Morales, J., Shulaev, V., Torres, M.A., and Mittler, R. (2011). Respiratory burst oxidases: the engines of ROS signaling. Curr Opin Plant Biol 14, 691–699.

Tanaka, H., Osakabe, Y., Katsura, S., Mizuno, S., Maruyama, K., Kusakabe, K., Mizoi, J., Shinozaki, K., and Yamaguchi-Shinozaki, K. (2012). Abiotic stress-inducible receptor-like kinases negatively control ABA signaling in Arabidopsis. Plant J 70, 599–613.

Torres, M.A., Dangl, J.L., and Jones, J.D. (2002). Arabidopsis gp91^phox^ homologues *AtrbohD* and *AtrbohF* are required for accumulation of reactive oxygen intermediates in the plant defense response. Proc Natl Acad Sci USA 99, 517–522.

Toyota, M., Spencer, D., Sawai-Toyota, S., Jiaqi, W., Zhang, T., Koo, A.J., Howe, G.A., and Gilroy, S. (2018). Glutamate triggers long-distance, calcium-based plant defense signaling. Science 361, 1112–1115.

Trost, B., Kusalik, A., and Napper, S. (2016). Computational analysis of the predicted evolutionary conservation of human phosphorylation sites. PLoS One 11, e0152809.

Vaattovaara, A., Brandt, B., Rajaraman, S., Safronov, O., Veidenberg, A., Luklová, M., Kangasjärvi, J., Löytynoja, A., Hothorn, M., Salojärvi, J., et al. (2019). Mechanistic insights into the evolution of DUF26-containing proteins in land plants. Commun Biol 2, 56.

Veidenberg, A., Medlar, A., and Löytynoja, A. (2016). Wasabi: An integrated platform for evolutionary sequence analysis and data visualization. Mol Biol Evol 33, 1126–1130.

Veronese, P., Nakagami, H., Bluhm, B., Abuqamar, S., Chen, X., Salmeron, J., Dietrich, R.A., Hirt, H., and Mengiste, T. (2006). The membrane-anchored *BOTRYTIS-INDUCED KINASE1* plays distinct roles in *Arabidopsis* resistance to necrotrophic and biotrophic pathogens. Plant Cell 18, 257–273.

Waszczak, C., Carmody, M., and Kangasjärvi, J. (2018). Reactive oxygen species in plant signaling. Annu Rev Plant Biol 69, 209–236.

Whalen, M.C., Innes, R.W., Bent, A.F., and Staskawicz, B.J. (1991). Identification of *Pseudomonas syringae* pathogens of *Arabidopsis* and a bacterial locus determining avirulence on both *Arabidopsis* and soybean. Plant Cell 3, 49–59.

Wrzaczek, M., Brosché, M., Salojärvi, J., Kangasjärvi, S., Idänheimo, N., Mersmann, S., Robatzek, S., Karpinski, S., Karpinska, B., and Kangasjärvi, J. (2010). Transcriptional regulation of the CRK/DUF26 group of receptor-like protein kinases by ozone and plant hormones in Arabidopsis. BMC Plant Biol 10, 95.

Wrzaczek, M., Rozhon, W., and Jonak, C. (2007). A Proteasome-regulated glycogen synthase kinase-3 modulates disease response in plants. J Biol Chem 282, 5249–5255.

Yadeta, K.A., Elmore, J.M., Creer, A.Y., Feng, B., Franco, J.Y., Rufian, J.S., He, P., Phinney, B., and Coaker, G. (2017). A cysteine-rich protein kinase associates with a membrane immune complex and the cysteine residues are required for cell death. Plant Physiol 173, 771–787.

Yao, J., Withers, J., and He, S.Y. (2013). Pseudomonas syringae infection assays in Arabidopsis. Methods Mol Biol 1011, 63–81.

Yeh, Y.H., Chang, Y.H., Huang, P.Y., Huang, J.B., and Zimmerli, L. (2015). Enhanced Arabidopsis pattern-triggered immunity by overexpression of cysteine-rich receptor-like kinases. Front Plant Sci 6, 322.

Yeh, Y.H., Panzeri, D., Kadota, Y., Huang, Y.C., Huang, P.Y., Tao, C.N., Roux, M., Chien, H.C., Chin, T.C., Chu, P.W., et al. (2016). The Arabidopsis Malectin-Like/LRR-RLK IOS1 Is Critical for BAK1-Dependent and BAK1-Independent Pattern-Triggered Immunity. Plant Cell 28, 1701–1721.

Yun, B.W., Feechan, A., Yin, M., Saidi, N.B., Le Bihan, T., Yu, M., Moore, J.W., Kang, J.G., Kwon, E., Spoel, S.H., et al. (2011). S-nitrosylation of NADPH oxidase regulates cell death in plant immunity. Nature 478, 264–268.

Zhang, J., Shao, F., Li, Y., Cui, H., Chen, L., Li, H., Zou, Y., Long, C., Lan, L., Chai, J., et al. (2007). A Pseudomonas syringae effector inactivates MAPKs to suppress PAMP-induced immunity in plants. Cell Host Microbe 1, 175–185.

Zhang, M., Chiang, Y.H., Toruno, T.Y., Lee, D., Ma, M., Liang, X., Lal, N.K., Lemos, M., Lu, Y.J., Ma, S., et al. (2018). The MAP4 kinase SIK1 ensures robust extracellular ROS burst and antibacterial immunity in plants. Cell Host Microbe 24, 379–391.

Zipfel, C., Robatzek, S., Navarro, L., Oakeley, E.J., Jones, J.D., Felix, G., and Boller, T. (2004). Bacterial disease resistance in *Arabidopsis* through flagellin perception. Nature 428, 764–767.

