## Supplemental Figures 1-7 for "CRK2 and C-terminal phosphorylation of NADPH oxidase RBOHD regulate ROS production in Arabidopsis"

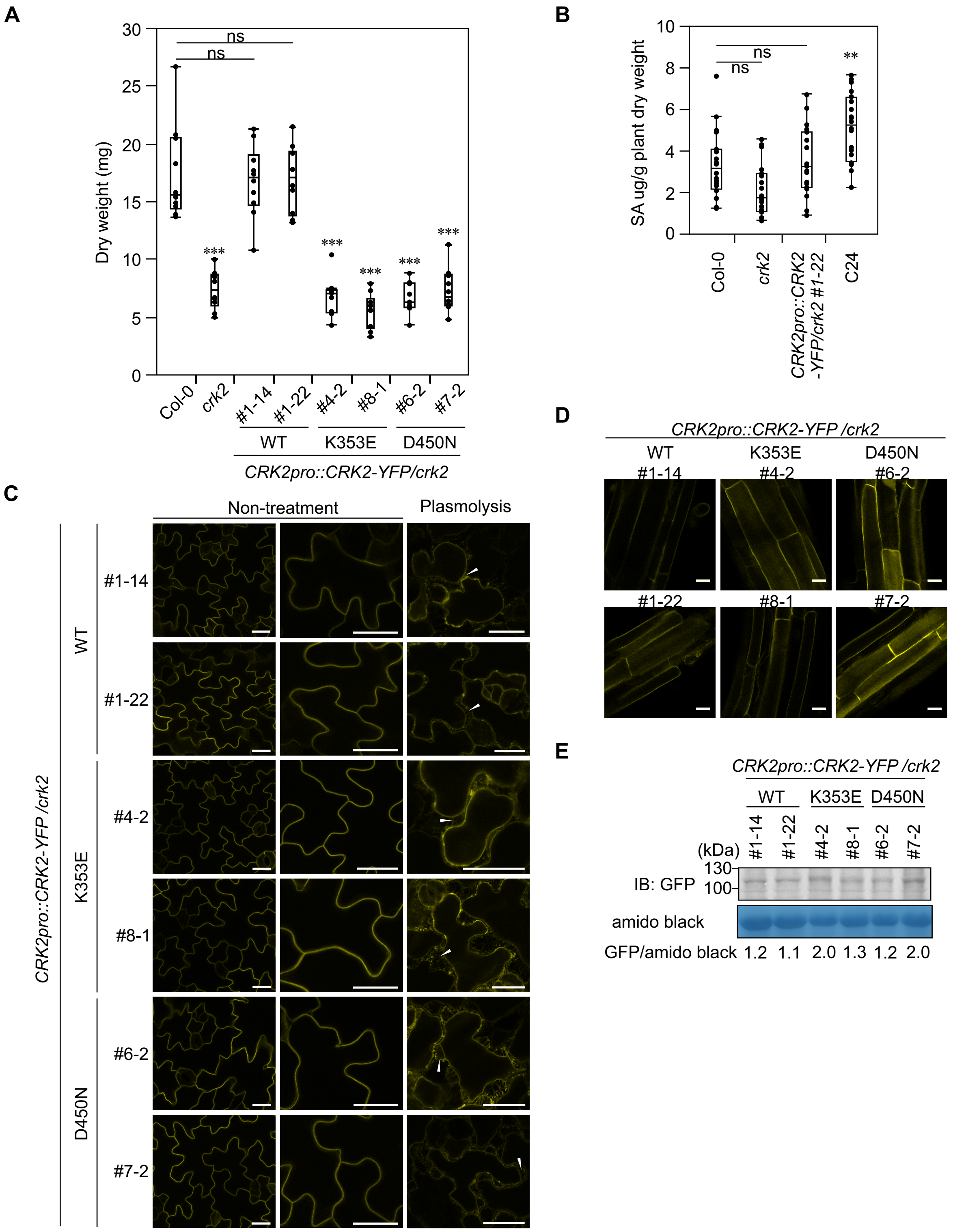


**Supplemental Figure 1. Complementation of *crk2* with *CRK2pro::CRK2-YFP* (Supports Figure 1).**

**(A)** Box plot shows dry weight of 21-day-old plants (n = 10). Differences compared with Col-0 were evaluated with One-way Anova with Tukey-Kramer HSD, *** p < 0.001, ns, not statistically significant. The experiment was repeated three times with similar results.

**(B)** Box plot shows salicylic acid accumulation level in Col-0, *crk2, CRK2pro::CRK2-YFP/crk2 #1-22* and C24 (n ≥ 19). 25-day-old plants were used. The experiment was repeated five times (three or four data points per experiment) and all data points are blotted. Differences compared with Col-0 were evaluated with One-way Anova with Tukey-Kramer HSD, ** p < 0.01, ns, not statistically significant.

(**C)** and **(D)** Subcellular localization of CRK2-YFP, CRK2$^{K353E}$-YFP and CRK2$^{D450N}$-YFP of 7-day-old seedlings.

**(C)** In leaves. Plasma membrane localization was confirmed using plasmolysis to visualize Hechtian strands (arrow head). Plasmolysis was induced by the application of 0.8 M mannitol. Scale bar = 25 µm.

**(D)** In roots. Scale bar = 20 µm.

**(E)** Expression of CRK2-YFP, CRK2$^{K353E}$-YFP and CRK2$^{D450N}$-YFP in 13-day-old seedlings were detected by anti-GFP antibody. Input proteins stained with amido black staining (lower panel). To quantify the expression level, GFP signal intensity was normalized by amido black signal intensity (GFP/amido black value).

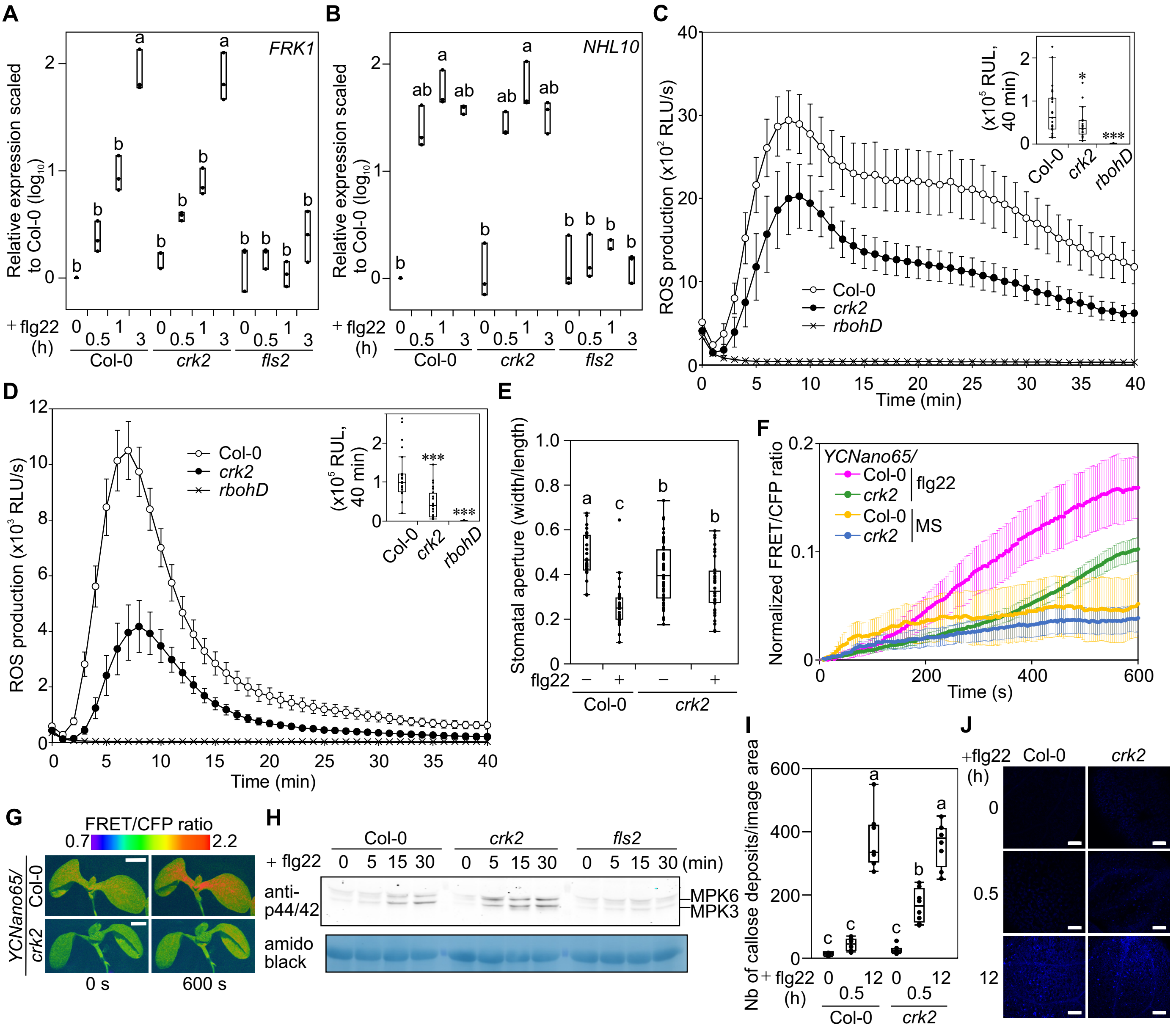


**Supplemental Figure 2. MAMP-triggered ROS production and molecular responses in *crk2* (Supports Figure 2).**

**(A)** and **(B)** Box plot shows quantitative real-time RT-PCR (qPCR) analysis of *FRK1* **(A)** or *NHL10* **(B)** transcripts in Col-0, *crk2* and *fls2* after treatment with flg22 (n = 3, biological replicates). 10-day-old plants were incubated in 1 μM flg22 solution and collected at indicated time (each data point contains 90 plants). Sample collection and qPCR were repeated three times with the same procedure and the three independent results were plotted. Transcript levels were calculated by comparison with non-treated Col-0 (Time = 0). Different letters indicate significant difference at $p < 0.05$ (One-way Anova with Tukey-Kramer HSD).

**(C)** and **(D)** Chitin- or AtPep1- induced ROS production in Col-0, *crk2* and *rbohD*. Leaf discs from 28-day-old plants were treated with 200 μg/mL chitin (**C**) or 1 μM AtPep1 (**D**). ROS production is expressed in relative luminescence units (RLU). Box plots show integration of ROS production for 40 min (upper right). Differences compared with Col-0 were evaluated with One-way Anova with Tukey-Kramer HSD. The experiment was repeated three times with similar results. **(C)** Values represent the mean ±SEM of n ≥ 21, * $p < 0.05$, *** $p < 0.001$. **(D)** Values represent the mean ±SEM of n = 24, *** $p < 0.001$.

**(E)** Quantification of flg22-induced stomatal closure in Col-0 (n = 28) and *crk2* (n ≥ 40). Leaf discs from 21-day-old plants were treated with 5 μM flg22 for 2 h. Letters indicate significant differences at $p < 0.05$ (One-way Anova with Tukey-Kramer HSD).

**(F)** Quantitative analysis of cytosolic $Ca^{2+}$ changes in response to 1×MS 1 % sucrose liquid media or 1 μM flg22 in 7-day-old *YCNano65* or *YCNano65/crk2* seedlings. Values represent the mean ±SEM of n = 9 (*YCNano65*) or n = 15 (*YCNano65/crk2*).

**(G)** Representative frame images of cytosolic $Ca^{2+}$ change in wild type and *crk2* plants. Bar = 0.5 mm.

**(H)** MAPK activation in Col-0, *crk2* and *fls2* in response to treatment with 1 μM flg22. 28-day-old plants (12 plants per genotype). Phosphorylated MPK3 and MPK6 were detected with anti-p44/42 MPK antibody (upper panel). Proteins stained with amido black staining (lower panel). The experiment was repeated three times with similar results.

**(I)** Quantification of flg22-induced callose deposition by aniline blue (n ≥ 8) in 7-day-old seedlings treatment with 10 μM flg22 for 30 min and 12 h. Letters indicate significant differences at $p < 0.05$ (One-way Anova with Tukey-Kramer HSD).

**(J)** Representative images of aniline blue stained leaves. Bar = 100 μm.

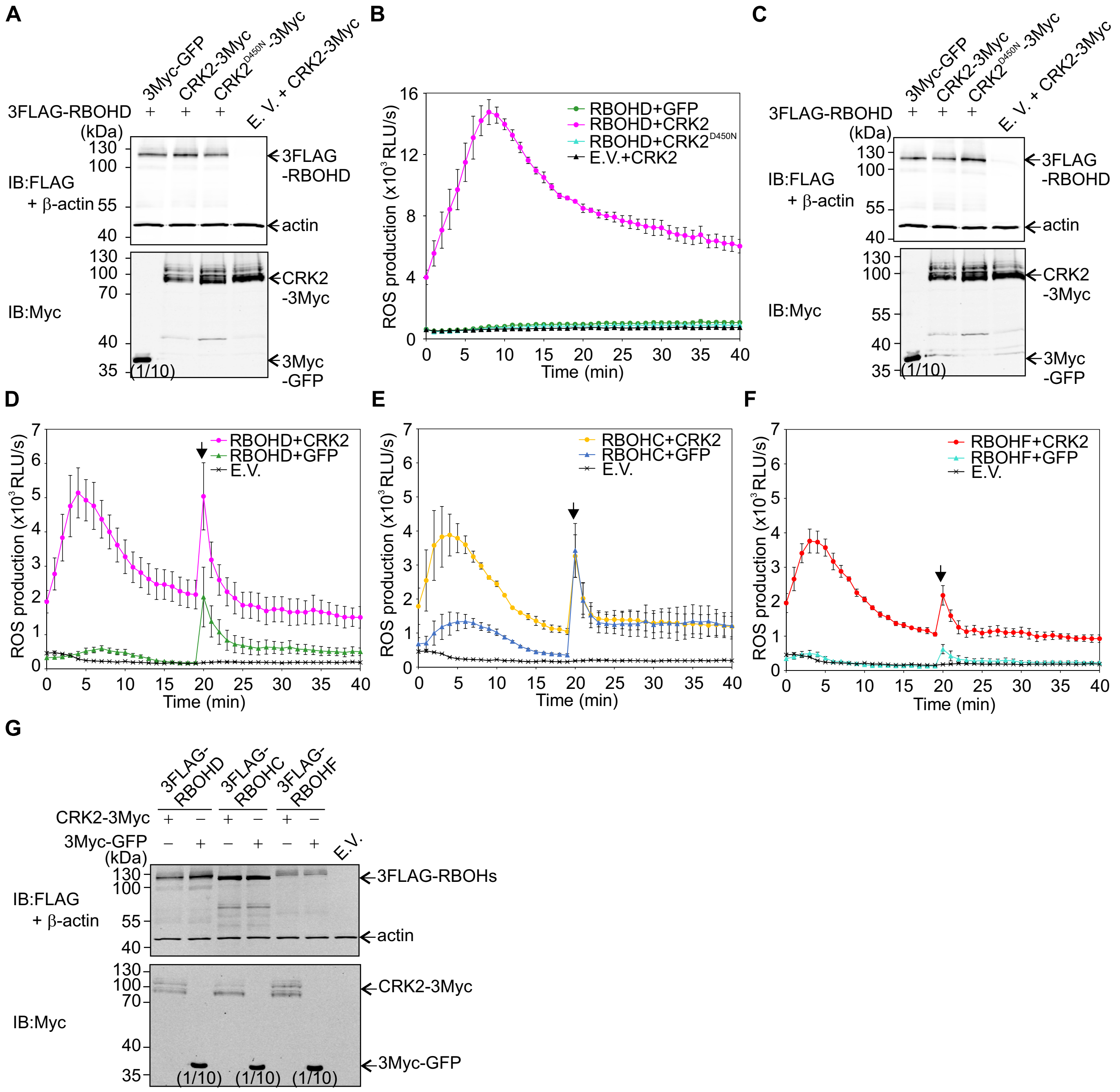


**Supplemental Figure 3. CRK2 modulates the ROS-producing activity of RBOHC, D and F in HEK293T cells (Supports Figure 3).**

**(A)** Expressed proteins were detected by anti-FLAG and anti-Myc antibodies (Figure 3A). 3FLAG-RBOHD: 107 kDa, CRK2-3Myc: 75.8 kDa, 3Myc-GFP: 31 kDa, b-actin 42 kDa. As a loading control, b-actin was used. Loading volume for anti-Myc antibody: 3FLAG-RBOHD + 3Myc-GFP (5 µL), the others (50 µL).

**(B)** ROS production in RBOHD-expressing HEK293T cells in $Ca^{2+}$-free buffer. 3FLAG-RBOHD was transiently co-expressed with either 3Myc-GFP or 3Myc-CRK2 (WT or D450N) in HEK293T cells. Values represent mean ±SEM of n = 3. E.V. = empty vector. The experiment was repeated three times with similar results.

**(C)** Proteins expressed in HEK293T cells were detected by anti-FLAG and anti-Myc antibodies (Supplemental Figure 3B). 3FLAG-RBOHD: 107 kDa, CRK2-3Myc: 75.8 kDa, 3Myc-GFP: 31 kDa, b-actin 42 kDa. As a loading control, b-actin was used. Loading volume for anti-Myc antibody: 3FLAG-RBOHD + 3Myc-GFP (5 µL), others (50 µL).

**(D)** to **(F)** ROS production of RBOHD-, RBOHC-, or RBOHF-expressing HEK293T cells. 3FLAG-RBOHD **(D)**, 3FLAG-RBOHC **(E)**, or 3FLAG-RBOHF **(F)** was transiently co-expressed with 3Myc-GFP or CRK2-3Myc in HEK293T cells, respectively. After 20 min of base line measurement, 1 µM ionomycin was added to the medium (black arrow). Values represent mean ±SEM of n = 3. The experiment was repeated two times with similar results.

**(G)** Proteins expressed in HEK293T cells were detected by anti-FLAG and anti-Myc antibodies (Supplemental Figures 3D to 3F). 3FLAG-RBOHD: 107 kDa, 3FLAG-RBOHC, 106 kDa, 3FLAG-RBOHF: 111 kDa, CRK2-3Myc: 75.8 kDa, 3Myc-GFP: 31 kDa, b-actin 42 kDa. As a loading control, b-actin was used. Loading volume for anti-Myc antibody: 3FLAG-RBOHs + 3Myc-GFP (5 µL), 3FLAG-RBOHs + CRK2-3Myc (50 µL).

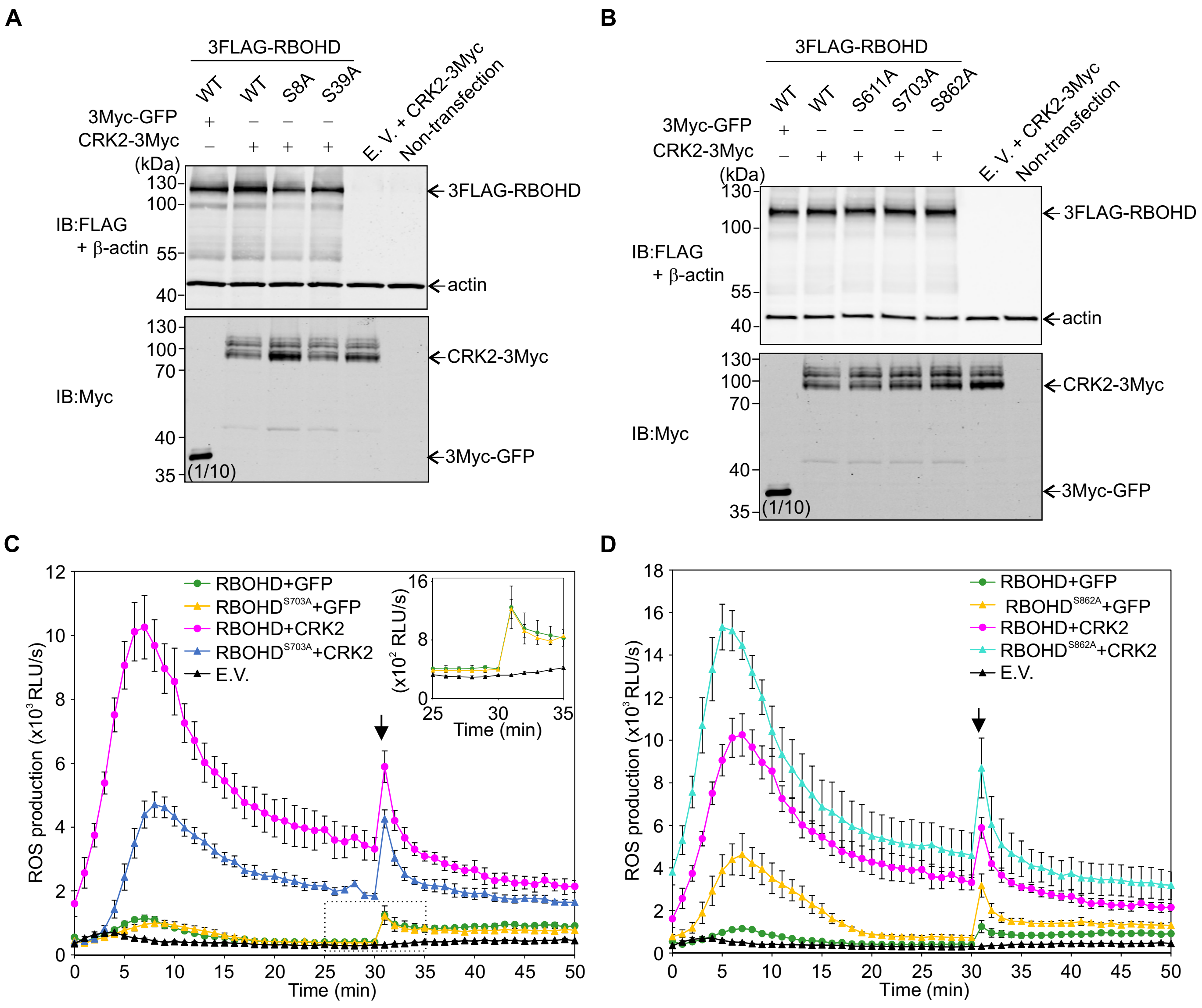


**Supplemental Figure 4. ROS production activity of RBOHD S703A and S862A in HEK293T cells (Supports Figure 5).**

**(A)** and **(B)** Proteins expressed in HEK293T cells were detected by anti-FLAG and anti-Myc antibodies (Figures 5A and 5B). 3FLAG-RBOHD: 107 kDa, CRK2-3Myc: 75.8 kDa, 3Myc-GFP: 31 kDa, b-actin 42 kDa. As a loading control, b-actin was used. Loading volume for anti-Myc antibody: 3FLAG-RBOHD + 3Myc-GFP (5 µL), others (50 µL).

**(A)** 3FLAG-RBOHD (WT, S8A or S39A) was transiently co-expressed with either 3Myc-GFP or CRK2-3Myc into HEK293T cells (Figure 5A).

**(B)** 3FLAG-RBOHD (WT, S611A, S703A or S862A) was transiently co-expressed with either 3Myc-GFP or CRK2-3Myc into HEK293T cells (Figure 5B).

**(C)** and **(D)** ROS production of RBOHD-expressing HEK293T cells. After 30 min 1 µM ionomycin was added to the medium (black arrow). Values represent mean ±SEM of n = 3. E.V. = empty vector. The experiment was repeated three times with similar results.

**(C)** 3FLAG-RBOHD (WT or S703A) was transiently co-expressed with either 3Myc-GFP or CRK2-3Myc into HEK293T cells. The right upper panel is the enlargement of ionomycin-induced transient ROS production of 3FLAG-RBOHD (WT or S703A) and 3Myc-GFP co-expressing cells (dashed box).

**(D)** 3FLAG-RBOHD (WT or S862A) was transiently co-expressed with either 3Myc-GFP or CRK2-3Myc into HEK293T cells.

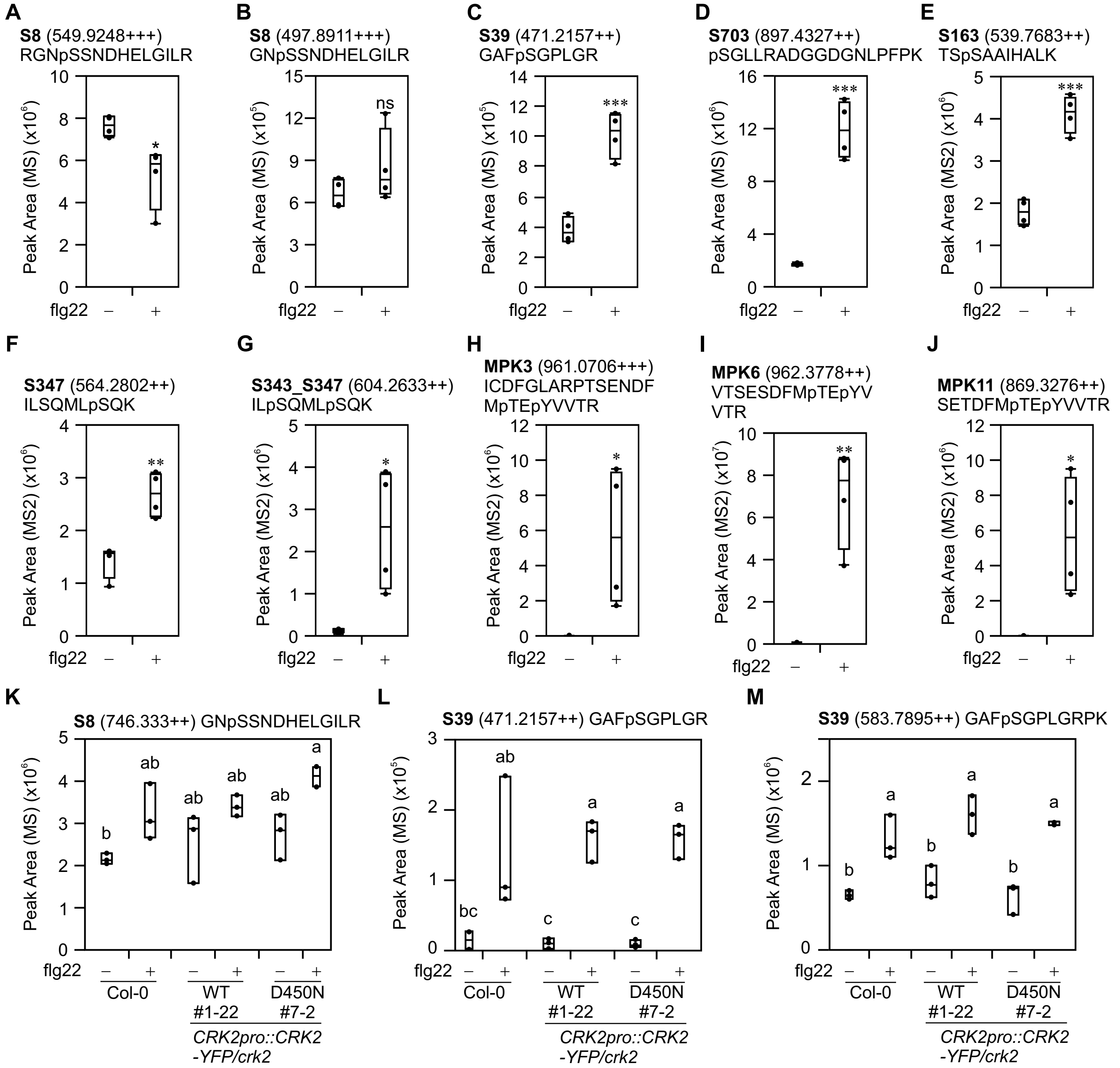


**Supplemental Figure 5. Quantification of RBOHD and MPK phosphorylation upon flg22 treatment (Supports Figure 6).**

**(A)** to **(J)** 12-day-old Col-0 seedlings were treated with water (-) or 1 µM flg22 (+) for 5 min. Total proteins were digested by trypsin for peptides from RBOHD N-terminal region and MPKs, by Lys-C for RBOHD S703 peptide. Peptides were enriched, and then selected phosphopeptides were quantified by LC-MS/MS. Box plots show MS2 fragment peak ion areas of indicated phosphopeptides (n = 4). Differences between water- or flg22-treated samples were evaluated with One-way Anova with Tukey-Kramer HSD, * P<0.05, ** P<0.01, *** p<0.001, ns, not statistically significant.

**(A)** and **(B)** RBOHD S8 residue. **(C)** RBOHD S39 residue. **(D)** RBOHD S703 residue. **(E)** RBOHD S163 residue. **(F)** RBOHD S347 residue. **(G)** RBOHD S343 and S347 residues. **(H)** MPK3 TEY motif. **(I)** MPK6 TEY motif. **(J)** MPK11 TEY motif.

**(K)** to **(M)** Quantification of RBOHD S8 and S39 phosphorylation in Col-0, *CRK2pro::CRK2-YFP/crk2* #1-22 and *CRK2pro::CRK2*$^{D450N}$*-YFP/crk2* #7-2 upon flg22 treatment. Leaf discs from 28-day-old plants were treated with water (-) or 200 nM flg22 (+) for 5 min. Total proteins were digested with trypsin and phosphopeptides were enriched, and then selected phosphopeptides were quantified by LC-MS/MS. Box plots show MS2 fragment peak ion areas of indicated phosphopeptides. Each data point contains 14 leaf discs. Letters indicate significant differences at p < 0.05 (One-way Anova with Tukey-Kramer HSD).

(**K**) RBOHD S8 residue (n ≥ 2).

(**L**) and (**M**) RBOHD S39 residue. n ≥ 2 (**L**), n = 3 (**M**).

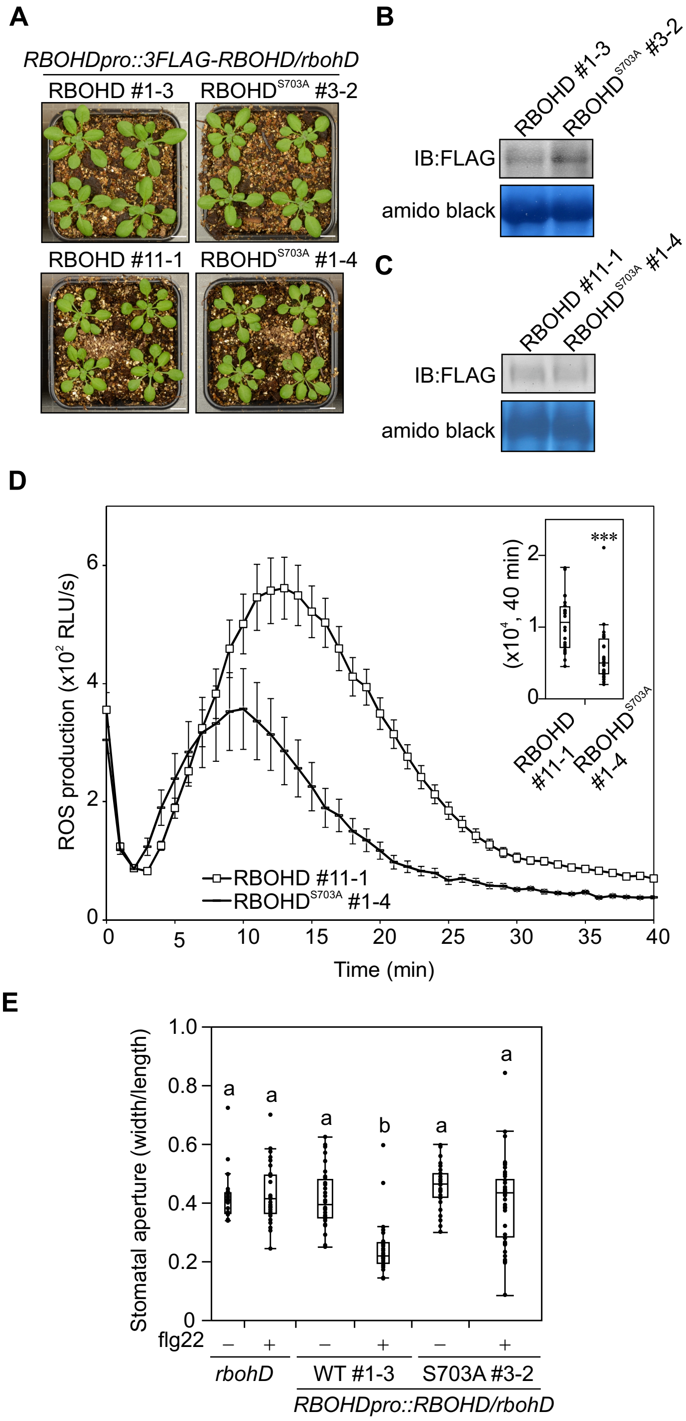


**Supplemental Figure 6. RBOHD S703 is involved in regulation of flg22-induced ROS production (Supports Figure 6).**

**(A)** Representative pictures of 21-day-old plants of *RBOHDpro::3FLAG-RBOHD/rbohD* #1-3, #11-1 and *RBOHDpro::3FLAG-RBOHD$^{S703A}$/rbohD* #3-2, #1-4 plants. Bar = 1 cm.

**(B)** and **(C)** Expressed proteins in 14-day-old *RBOHDpro::3FLAG-RBOHD/rbohD* and *RBOHDpro::3FLAG-RBOHD$^{S703A}$/rbohD*. 3FLAG-RBOHD was detected by anti-FLAG antibody. Input proteins stained with amido black staining (lower panel).

**(B)** *RBOHDpro::3FLAG-RBOHD/rbohD* #1-3 and *RBOHDpro::3FLAG-RBOHD$^{S703A}$/rbohD* #3-2.

**(C)** *RBOHDpro::3FLAG-RBOHD/rbohD* #11-1 and *RBOHDpro::3FLAG-RBOHD$^{S703A}$/rbohD* #1-4.

**(D)** flg22-induced ROS production in *RBOHDpro::3FLAG-RBOHD/rbohD* #11-1 and *RBOHDpro::3FLAG-RBOHD$^{S703A}$/rbohD* #1-4. Leaf discs from 28-day-old plants were treated with 200 nM flg22. Box plots show integration of ROS production for 40 min (upper right). Values represent the mean ±SEM of n = 24. Difference between lines was evaluated with One-way Anova with Tukey-Kramer HSD, *** $p < 0.001$.

**(E)** Quantification of flg22-induced stomatal closure in *rbohD* (n ≥ 21), *RBOHDpro::3FLAG-RBOHD/rbohD* #1-3 (n ≥ 35) and *RBOHDpro::3FLAG-RBOHD$^{S703A}$/rbohD* #3-2 (n ≥ 35). Leaf discs from 21-day old plants were treated with 5 µM flg22 for 2 h. Letters indicate significant differences at $p < 0.05$ (One-way Anova with Tukey-Kramer HSD).

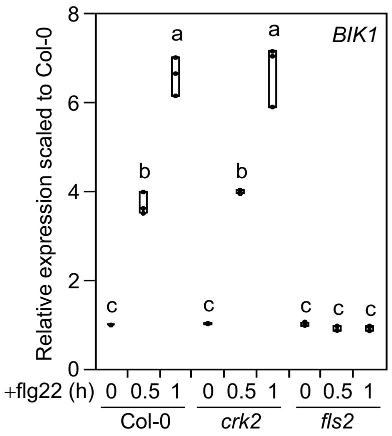


**Supplemental Figure 7. Reduced ROS production in *crk2* is not due to lower expression of *BIK1* (Supports Figure 2).**

Box plot shows quantitative real-time RT-PCR (qPCR) analysis of *BIK1* transcripts in Col-0, *crk2* and *fls2* after treatment with flg22 (n = 3, biological replicates). 10-day-old plants were incubated in 1 μM flg22 solution and collected at indicated time (each data point contains 90 plants). Sample collection and qPCR were repeated three times with the same procedure and the three independent results were plotted. Transcript levels were calculated by comparison with non-treated Col-0 (Time = 0). Different letters indicate significant difference at $p < 0.05$ (One-way Anova with Tukey-Kramer HSD).
