## Supplemental Tables 1-5 for "CRK2 and C-terminal phosphorylation of NADPH oxidase RBOHD regulate ROS production in Arabidopsis"

**Supplemental Table 1.** ***In vitro*** **phosphorylation sites of 6His-MBP-RBOHDcyto by 6His-GST-CRK2cyto**

Trypsin

| Phosphorylation site | Peptide Sequence | Best Site Probabilities | Mascot Ions Score | Replicates |
|---|---|---|---|---|
| S8 | GNpSSNDHELGILR | 100 | 97 | 5 |
| S8 | RGNpSSNDHELGILR | 100 | 63.61 | 3 |
| S39 | GAFpSGPLGRPK | 100 | 48 | 5 |
| S611 | AFRpSSIKPVK | 99.69 | 30 | 5 |
| S611 | pSSIKPVK | 50 | 35 | 4 |
| S703 | TVFSEVCKPPTAGKpSGLLR | 100 | 83.29 | 6 |
| S703 | pSGLLR | 100 | 43 | 2 |
| S862 | VKpSHFAKPNWR | 100 | 45 | 5 |

Lys-C

| Phosphorylation site | Peptide Sequence | Best Site Probabilities | Mascot Ions Score | Replicates |
|---|---|---|---|---|
| S703 | pSGLLRADGGDGNLPFPK | 100 | 29 | 1 |

The 6His-MBP-RBOHD cytosolic regions were incubated with 6His-GST-CRK2cyto. The 6His-MBP-RBOHDcyto bands were excised from a SDS polyacrylamide gel and subsequently digested by trypsin or Lys-C. The peptides were analyzed by LC-MS/MS. Phosphorylated peptides are designated as pS.

**Supplemental Table 2. The numbers of Serine (S) and Threonine (T) residues in C-terminus of NADPH oxidases in plants and animals.**

| Taxon | Found (ST) | Found (S) | Found (T) |
|---|---|---|---|
| NOX5_HUMAN_beta | 40 | 25 | 15 |
| NOX2_HUMAN | 33 | 17 | 16 |
| AT5G47910 | 35 | 21 | 14 |
| Carubv10007974m.g | 36 | 20 | 16 |
| Prupe.1G211000 | 44 | 29 | 15 |
| Aqcoe5G319800.1 | 35 | 21 | 14 |
| Solyc03g117980.2 | 39 | 26 | 13 |
| Solyc06g068680.2 | 39 | 23 | 16 |
| Prupe.5G138300 | 38 | 26 | 12 |
| AT4G25090 | 40 | 21 | 19 |
| Carubv10006425m.g | 44 | 24 | 20 |
| Carubv10004156m.g | 40 | 24 | 16 |
| AT5G51060 | 36 | 22 | 14 |
| Carubv10028541m.g | 36 | 21 | 15 |
| AT5G07390 | 48 | 32 | 16 |
| ATR0582G212 | 42 | 24 | 18 |
| LOC_Os11g33120 | 41 | 25 | 16 |
| Sobic.005G139700 | 41 | 25 | 16 |
| AT1G09090 | 45 | 27 | 18 |
| Carubv10008289m.g | 43 | 26 | 17 |
| Solyc01g099620.2 | 41 | 24 | 17 |
| Prupe.6G321500 | 41 | 23 | 18 |
| Aqcoe5G083800.1 | 38 | 22 | 16 |
| LOC_Os01g25820.1 | 43 | 26 | 17 |
| Sobic.003G161500.3 | 38 | 23 | 15 |
| LOC_Os12g35610 | 40 | 21 | 19 |
| Sobic.008G118700.1 | 43 | 26 | 17 |
| ATR0681G031 | 52 | 31 | 21 |
| AT1G19230 | 56 | 32 | 24 |
| Solyc07g042460.1 | 58 | 35 | 23 |
| Solyc05g025690_80_70.1 | 52 | 31 | 21 |
| Aqcoe7G437300.1 | 59 | 33 | 26 |
| Prupe.7G193000 | 55 | 31 | 24 |
| LOC_Os08g35210 | 50 | 29 | 21 |
| Sobic.007G148300.1 | 45 | 25 | 20 |
| LOC_Os09g26660 | 52 | 29 | 23 |
| Sobic.002G214200.1 | 51 | 30 | 21 |
| AT1G64060 | 47 | 28 | 19 |
| Carubv10021686m.g | 48 | 28 | 20 |
| Solyc08g081690.2 | 45 | 26 | 19 |

| | | | |
|---|---|---|---|
| Prupe.5G107400 | 46 | 26 | 20 |
| Aqcoe5G438000.1 | 40 | 23 | 17 |
| LOC_Os01g53294 | 43 | 23 | 20 |
| Sobic.003G287400.1 | 42 | 23 | 19 |
| LOC_Os05g45210 | 42 | 25 | 17 |
| Sobic.009G206500.1 | 43 | 25 | 18 |
| AT4G11230 | 52 | 33 | 19 |
| Carubv10003524m.g | 52 | 35 | 17 |
| ATR0691G180 | 35 | 19 | 16 |
| Mapoly0258s0001 | 49 | 33 | 16 |
| Mapoly0046s0097 | 45 | 24 | 21 |
| ATR0730G224 | 42 | 27 | 15 |
| Aqcoe3G372000.1 | 44 | 25 | 19 |
| Prupe.5G204900 | 41 | 27 | 14 |
| Sobic.001G303600 | 36 | 25 | 11 |
| ATR0594G041 | 47 | 24 | 23 |
| ATR0743G058 | 45 | 20 | 25 |
| LOC_Os01g61880 | 46 | 26 | 20 |
| Sobic.003G347520.1 | 47 | 26 | 21 |
| LOC_Os05g38980 | 49 | 28 | 21 |
| Sobic.003G175000.1 | 43 | 25 | 18 |
| AT3G45810 | 50 | 35 | 15 |
| Carubv10019407m.g | 47 | 31 | 16 |
| AT5G60010 | 44 | 29 | 15 |
| Carubv10028111m.g | 43 | 29 | 14 |
| Aqcoe2G090100.1 | 42 | 22 | 20 |
| Solyc06g075570.1 | 45 | 29 | 16 |
| Solyc11g072800.1 | 39 | 22 | 17 |
| Prupe.6G088800 | 49 | 30 | 19 |
| Minimum | 33 | 17 | 11 |
| Maximum | 59 | 35 | 26 |

**Supplemental Table 3. Progeny of *CRK2/crk2 BIK1/bik1* parent and *CRK2/crk2 bik1/bik1* parent**

| Parent genotype | | Progeny genotype | | Observed | | Expected |
|---|---|---|---|---|---|---|
| **F1** | | **F2** | | | | |
| *CRK2/crk2* | *BIK1/bik1* | *CRK2/CRK2* | *BIK1/BIK1* | | 20 | 12.4375 |
| | | *CRK2/CRK2* | *BIK1/bik1* | | 29 | 24.875 |
| | | *CRK2/CRK2* | *bik1/bik1* | | 10 | 12.4375 |
| | | *CRK2/crk2* | *BIK1/BIK1* | | 33 | 24.875 |
| | | *CRK2/crk2* | *BIK1/bik1* | | 75 | 49.75 |
| | | *CRK2/crk2* | *bik1/bik1* | | 26 | 24.875 |
| | | *crk2/crk2* | *BIK1/BIK1* | | 6 | 12.4375 |
| | | *crk2/crk2* | *BIK1/bik1* | | 0 | 24.875 |
| | | *crk2/crk2* | *bik1/bik1* | | 0 | 12.4375 |
| | | | | Total | 199 | P < 0.001 |
| **F2** | | **F3** | | | | |
| *CRK2/crk2* | *bik1/bik1* | *CRK2/CRK2* | *bik1/bik1* | | 9 | 16.5 |
| | | *CRK2/crk2* | *bik1/bik1* | | 57 | 33 |
| | | *crk2/crk2* | *bik1/bik1* | | 0 | 16.5 |
| | | | | Total | 66 | P < 0.001 |

The genotypes of F2 and F3 progenies were determined by PCR. Observed, the number of individuals observed; Expected, the expected number based on Mendelian inheritance. Chi-square test was used to determine the probability (P) of which the deviation of the observed value from the expected value was due to chance.

**Supplemental Table 4. Primer sequences**

| Primer | Sequence (5'-3') | |
|---|---|---|
| CRK2-for-pTOPO | CACCATGAAGAAAGAACCTGTCCATATC | pENTR /D-TOPO |
| CRK2ns-rev | TCTACCATAAAAGGAACTTTG | |
| B4-RBOHDpro for | GGGGACAACTTTGTATAGAAAAGTTGCCCCCCTCTAGTTCTTGT GATGC | pDONRP4 P1R/zeo |
| B1-RBOHDpro rev | GGGGACTGCTTTTTTGTACAAACTTGGCGAATTCGAGAAACCAA AAAGATCTC | |
| B1-3FLAG-RBOHD for | GGGGACAAGTTTGTACAAAAAAGCAGGCTGTATGGACTACAAAG ACCACGATG | pDONR /zeo |
| B2-3FLAG-RBOHD rev | GGGGACCACTTTGTACAAGAAAGCTGGGTTCTAGAAGTTCTCTT TGTGGAAGTC | |
| pOPIN-CRK2cyto for | AAGTTCTGTTTCAGGGCCCGAAGAGGAAGAGAAGAGGATC | pOPINK |
| pOPIN-CRK2cyto rev | ATGGTCTAGAAAGCTTTATCTACCATAAAAGGAACTTTGTGA | |
| pOPIN-RBOHD/C for | AAGTTCTGTTTCAGGGCCCGATGCTCCGTGCTTTCAGATCAAGC | pOPINM |
| pOPIN-RBOHD/C rev | ATGGTCTAGAAAGCTTTACTAGAAGTTCTCTTTGTGGAAGTC | |
| pOPIN-RBOHD/C1 rev | ATGGTCTAGAAAGCTTTAGAGGAGTACCACGTCGTATTTC | |
| pOPIN-RBOHD/C2 for | AAGTTCTGTTTCAGGGCCCGAAACCTCCTACCGCCGGTAAAAGC | |
| pOPIN-RBOHD/C2 rev | ATGGTCTAGAAAGCTTTATTCCTCGTACACACTCGTGC | |
| pOPIN-RBOHD/C3 for | AAGTTCTGTTTCAGGGCCCGAAAGCTTATTTCTACTGGGTGAC | |
| KpnI-MCS-3Myc-stop-XbaI for | GGTACCGGATCCGGCGGCCGCGAACAAAAGTTGATTTCAGAAG AAGATCTGGAACAAAAGTTGATTTCAGAAGAAGATCTGGAACAA AAGTTGATTTCAGAAGAAGATCTGTGATCTAGA | pEF1/myc-His B |
| KpnI-MCS-3Myc-stop-XbaI rev | TCTAGATCACAGATCTTCTTCTGAAATCAACTTTTGTTCCAGATCT TCTTCTGAAATCAACTTTTGTTCCAGATCTTCTTCTGAAATCAACT TTTGTTCGCGGCCGCCGGATCCGGTACC | |
| BamHI-koz-CRK2op for | TTTTGGATCCGCCGCCACCATGAAGAAG | pEF1-MCS-3Myc |
| NotI-CRK2op rev | TTTTTTGCGGCCGCCCCTGCCGTAGAAG | |
| BamHI- RBOHD-for | CCGGATCCATGAAAATGAGACGAGGCAATTC | pcDNA3.1-3FLAG-MCS |
| BamHI-RBOHD rev | GCGGATCCCTAGAAGTTCTCTTTGTGGAAGTC | |
| RBOHD S8A rev | CATGGTCGTTACTAGCATTGCCTCGTCTC | mega-primer |
| RBOHD S39A rev | GCCAAGCGGACCAGCAAAGGCACCACGG | |
| RBOHD S611A rev | CGGTTTAATGCTAGCTCTGAAAGCACG | |
| RBOHD S703A rev | CTCGGAGAAGACCAGCTTTACCGGCGGTAG | |
| RBOHD S862A rev | GTTTAGCGAAGTGAGCCTTGACACGTGTAC | |
| CRK2 K353E rev | CCGCTCCACAGCGATATCTCTTCCGTC | |
| CRK2 D450N rev | TGCTTTTATGTTTCTGTGAATAATTTTC | |
| CRK2op K353E rev | GCGCTCCACGGCGATGTCTCTGCCATC | |
| CRK2op D450N rev | GGCCTTGATATTGCGGTGGATGAT | |
| CRK2 for | GCTAACTATGGTCTTGCGCAG | genotyping |
| CRK2 rev | CAAAGATGAATCGATCAAGGC | |
| BIK1 for | CAGGTCACTTGAATGCAAGAAGCG | |
| BIK1 rev | GGGTATGGGACATGTAACCGGAAA | |

| | | |
|---|---|---|
| LbaI | TGGTTCACGTAGTGGGCCATCG | |
| RbohD for | GTCGCCAAAGGAGGCGCCGA | |
| RbohD rev | GGATACTGATCATAGGCGTGGCTCCA | |
| dSpm1 | CTTATTTCAGTAAGAGTGTGGGGTTTTGG | |
| q-NHL10 for | TTCCTGTCCGTAACCCAAAC | qRT-PCR |
| q-NHL10 rev | CCCTCGTAGTAGGCATGAGC | |
| q-FRK1 for | CGGTCAGATTTCAACAGTTGTC | |
| q-FRK1 rev | AATAGCAGGTTGGCCTGTAATC | |
| q-BIK1 for | TGGGCTCGACCGTACCTCACA | |
| q-BIK1 rev | CGGGCGCGACTTGGGTTCAA | |
| q-YLS8 for | TTACTGTTTCGGTTGTTCTCCATTT | |
| q-YLS8 rev | CACTGAATCATGTTCGAAGCAAGT | |
| q-TIP41 for | GTGAAAACTGTTGGAGAGAAGCAA | |
| q-TIP41 rev | TCAACTGGATACCCTTTCGCA | |
| q-SAND for | CAGACAAGGCGATGGCGATA | |
| q-SAND rev | GCTTTCTCTCAAGGGTTTCTGGGT | |

**Supplemental Table 5. One-way ANOVA results**

| Figure | Label | Oneway Anova | | | |
|---|---|---|---|---|---|
| | | **F Ratio** | **df** | **Error df** | **p value** |
| 1 | (C) | 71.5559 | 7 | 71 | < 0.0001 |
| 2 | (A) | 9.2282 | 3 | 73 | < 0.0001 |
| | (B) | 8.8777 | 3 | 76 | < 0.0001 |
| | (C) | 566.5661 | 11 | 42 | < 0.0001 |
| 6 | (A) | 0.4745 | 1 | 6 | 0.5167 |
| | (B) | 51.3297 | 1 | 6 | 0.0004 |
| | (C) | 41.0851 | 1 | 6 | 0.0007 |
| | (D) | 63.5965 | 5 | 12 | < 0.0001 |
| | (E) | 4.4509 | 1 | 45 | 0.0405 |
| | (F) | 10.7342 | 4 | 25 | < 0.0001 |
| Supplemental 1 | (A) | 48.2539 | 7 | 22 | < 0.0001 |
| | (B) | 12.9687 | 3 | 75 | < 0.0001 |
| Supplemental 2 | (A) | 9.4471 | 11 | 24 | < 0.0001 |
| | (B) | 9.1059 | 11 | 24 | < 0.0001 |
| | (C) | 24.1435 | 2 | 65 | < 0.0001 |
| | (D) | 44.5132 | 2 | 69 | < 0.0001 |
| | (E) | 20.5466 | 3 | 140 | < 0.0001 |
| | (I) | 86.8011 | 5 | 43 | < 0.0001 |
| Supplemental 5 | (A) | 9.355 | 1 | 6 | 0.0223 |
| | (B) | 1.7274 | 1 | 6 | 0.2367 |
| | (C) | 51.1741 | 1 | 6 | 0.0003 |
| | (D) | 87.8835 | 1 | 6 | < 0.0001 |
| | (E) | 71.832 | 1 | 6 | 0.0001 |
| | (F) | 22.8032 | 1 | 6 | 0.0031 |
| | (G) | 10.9184 | 1 | 6 | 0.0163 |
| | (H) | 8.0906 | 1 | 6 | 0.0294 |
| | (I) | 33.9863 | 1 | 6 | 0.0011 |
| | (J) | 11.6362 | 1 | 6 | 0.0143 |
| | (K) | 4.2965 | 5 | 11 | 0.0206 |
| | (L) | 8.4091 | 5 | 11 | 0.0017 |
| | (M) | 17.8271 | 5 | 12 | < 0.0001 |
| Supplemental 6 | (D) | 15.4533 | 1 | 46 | 0.0003 |
| | (E) | 19.5685 | 5 | 196 | < 0.0001 |
| Supplemental 7 | - | 220.624 | 8 | 18 | < 0.0001 |
